# TCAIM controls effector T cell generation by preventing Mitochondria-Endoplasmic Reticulum Contact Site-initiated Cholesterol Biosynthesis

**DOI:** 10.1101/2021.04.20.440500

**Authors:** Christina Iwert, Julia Stein, Christine Appelt, Katrin Vogt, Roman Josef Rainer, Katja Tummler, Kerstin Mühle, Katarina Stanko, Julia Schumann, Doreen Uebe, Karsten Jürchott, Jan Lisec, Katharina Janek, Christoph Gille, Kathrin Textoris-Taube, Somesh Sai, Ansgar Petersen, Anja A. Kühl, Edda Klipp, Christian Meisel, Birgit Sawitzki

## Abstract

T cells need to adapt their cellular metabolism for effector cell differentiation. This relies on alterations in mitochondrial physiology. Which signals and molecules regulate those alterations remains unclear. We recently reported, that the mitochondrial protein TCAIM inhibits activation-induced changes in mitochondrial morphology and function and thus, CD4^+^ effector T cell formation. Using conditional TCAIM knock-in (KI) and knockout (KO) mice, we now show that it also applies to CD8^+^ T cells and more importantly, delineate the molecular processes in mitochondria by which TCAIM controls effector cell differentiation. TCAIM KI resulted in reduced activation-induced HIF1α protein expression. Metabolomics and transcriptional data in combination with mathematical flux modeling revealed an impaired induction of anabolic pathways, especially of the mevalonate pathway and cholesterol biosynthesis in TCAIM KI CD8^+^ T cells. Addition of cholesterol completely rescued HIF1α protein expression, activation and proliferation of TCAIM KI CD8^+^ T cells. At the molecular level, TCAIM delayed activation-induced mitochondria-ER contact (MERC) formation by binding to MERC promoting proteins such as RMD3 and VDAC2. In summary, we demonstrate that TCAIM suppresses effector cell differentiation by inhibiting MERC formation, which induce HIF1α-mediated increase in cellular metabolism and cholesterol biosynthesis.

## Introduction

In order to cope with increased metabolic demands arising with T cell activation and effector differentiation, T cells need to upregulate catabolic but also to induce anabolic processes. Upregulation of catabolic processes mainly comprises elevations of aerobic glycolysis, glutaminolysis and oxidative phosphorylation (OXPHOS) necessary for profound building block(s) and adenosine triphosphate (ATP) synthesis. Newly generated building blocks are then funneled into anabolic pathways to enable nucleotide, protein and lipid synthesis to ensure DNA replication, cell growth and proliferation as well as the acquisition of effector functions^1^.

Especially, effector T cell differentiation and function relies on upregulation of aerobic glycolysis^2^, which is regulated by the two transcription factors MYC and HIF1α. Whereas MYC is essential to initiate aerobic glycolysis during T cell activation^3^, HIF1α sustains aerobic glycolysis throughout T cell activation to enforce effector T cell differentiation and, more importantly, effector function^4^. HIF1α signaling has been shown to be indispensable for Granzyme B expression, CD62L downregulation and CD8^+^ T cell migration^4, 5^. Therefore, HIF1α activation is crucial for effector T cell function and recruitment to the inflammatory environment.

Metabolic adaptations necessary for effector T cell differentiation have been shown to depend on alterations in mitochondrial physiology^6^. OXPHOS is strongly upregulated upon T cell activation, which is accompanied by an increase in mitochondrial membrane potential (ΔΨm) and mitochondrial reactive oxygen species (mROS) generation^7^. Whereas an increase in ΔΨm is associated with an enhanced effector differentiation and, more importantly, effector cytokine secretion in T cells^8^, mROS have been shown to act as redox signaling molecules regulating gene expression and proliferation^9–11^. Besides mROS, mitochondria also actively shape cellular transcription through the release of TCA cycle derived metabolites^12^. Moreover, effector T cell differentiation has been linked to mitochondrial fission and cristae resolution as a prerequisite for acquisition of aerobic glycolysis^13^. Failure to engage either will result in deregulated T cell activation and differentiation^14^. Thus, mitochondria function as central immunometabolic hubs to guide T cell activation and differentiation. However, which signals and molecules regulate those activation-induced alterations in mitochondrial physiology still remains largely unclear.

We recently reported that the mitochondrial protein T cell activation inhibitor, mitochondrial (TCAIM) is downregulated upon T cell activation. Notably, reinforced TCAIM expression inhibits effector T cell differentiation of conventional CD4^+^ T cells leading to full acceptance of allogeneic skin grafts^15^. TCAIM exclusively locates within mitochondria^16^ where it might interfere with activation-induced changes in mitochondrial morphology and function^15^.

Here, we hypothesized that TCAIM acts as an important link between activation-induced mitochondrial alterations and changes in cellular metabolism driving full effector cell differentiation. To investigate this, we utilized conditional TCAIM KI and KO mice, overexpressing TCAIM or displaying a loss of function in the TCAIM protein, to examine which molecular processes in mitochondria control CD8^+^ T cell effector differentiation. We found that TCAIM KI strongly impaired CD8^+^ T cell proliferation and effector differentiation as well as upregulation of HIF1α protein expression and transcription of genes encoding for enzymes of the mevalonate pathway and cholesterol biosynthesis. Importantly, addition of cholesterol rescued T cell activation and proliferation of TCAIM KI CD8^+^ T cells. At the molecular level, we showed that TCAIM delays activation-induced interaction between mitochondria and the endoplasmic reticulum (ER) by binding to voltage-dependent anion-selective channel 2 (VDAC2) and regulator of microtubule dynamics protein 3 (RMD3), both of which promote mitochondria-ER contact (MERC) formation. In summary, we provide ce that TCAIM negatively regulates the activation-induced increase in MERC ion, thereby, preventing the upregulation of anabolic processes, particularly the onate pathway and cholesterol biosynthesis, thus, resulting in impaired COW T cell ion and proliferation.

## Results

### TCAIM KI CD8^+^ T cells show reduced glycolytic flux and biosynthesis as well as impaired CD8^+^ T cell differentiation

Upregulation of anabolic processes is a hallmark of T cell activation and differentiation. To investigate whether TCAIM interferes with CD8^+^ T cell anabolic upregulation upon activation, we performed a non-targeted metabolite analysis. For this, we used mice with a conditional T cell specific Tcaim gene overexpression (KI) and Cd4-Cre wild type (wt) controls from littermates or parallel breeding. Isolated naïve CD8^+^ T cells from spleen and lymph nodes of these mice were polyclonally activated in vitro with αCD3 and αCD28 antibodies for 60 h.

### Metabolite levels were measured using gas chromatography mass spectrometry (GC/APCI-MS)

Comparing the levels of a chosen fraction of detected metabolites from TCAIM KI CD8^+^ T cells with that of wt CD8^+^ T cells we found, that TCAIM KI CD8^+^ T cells showed increased glucose levels, while intermediates and especially end products of the glycolytic pathway like pyruvate or lactic acid were reduced suggesting a diminished glycolytic flux due to TCAIM KI (Fig. 1a upper row and Extended Data Fig. 1a). Also, intermediates of the phospholipid biosynthesis pathway like glycerol-3-phosphate or sphingosine as well as levels of non-essential amino acids derived either from glycolysis or TCA cycle intermediates e.g. alanine and serine or proline and aspartate, respectively, were reduced (Fig. 1a middle and lower row). In contrast, citrate and α-ketoglutarate early intermediates of the TCA cycle were increased while late TCA cycle intermediates like malate or fumarate were strongly decreased (Extended Data Fig. 1b). Additionally, glutamine, starting point of the glutaminolysis pathway, was enriched while glutamate showed similar levels (Extended Data Fig. 1c) suggesting a reduced support of the TCA cycle through a diminished glutamine breakdown. Interestingly, although reduced glycolysis and TCA cycle end products or lower levels of biosynthesis related metabolites suggest an overall diminished metabolism in TCAIM KI CD8^+^ T cells, intermediates of the pentose phosphate pathway as ribose-5-phosphate and other sugar metabolites such as fructose were enriched in TCAIM KI CD8^+^ T cells (Extended Data Fig. 1d).

**Fig. 1.**
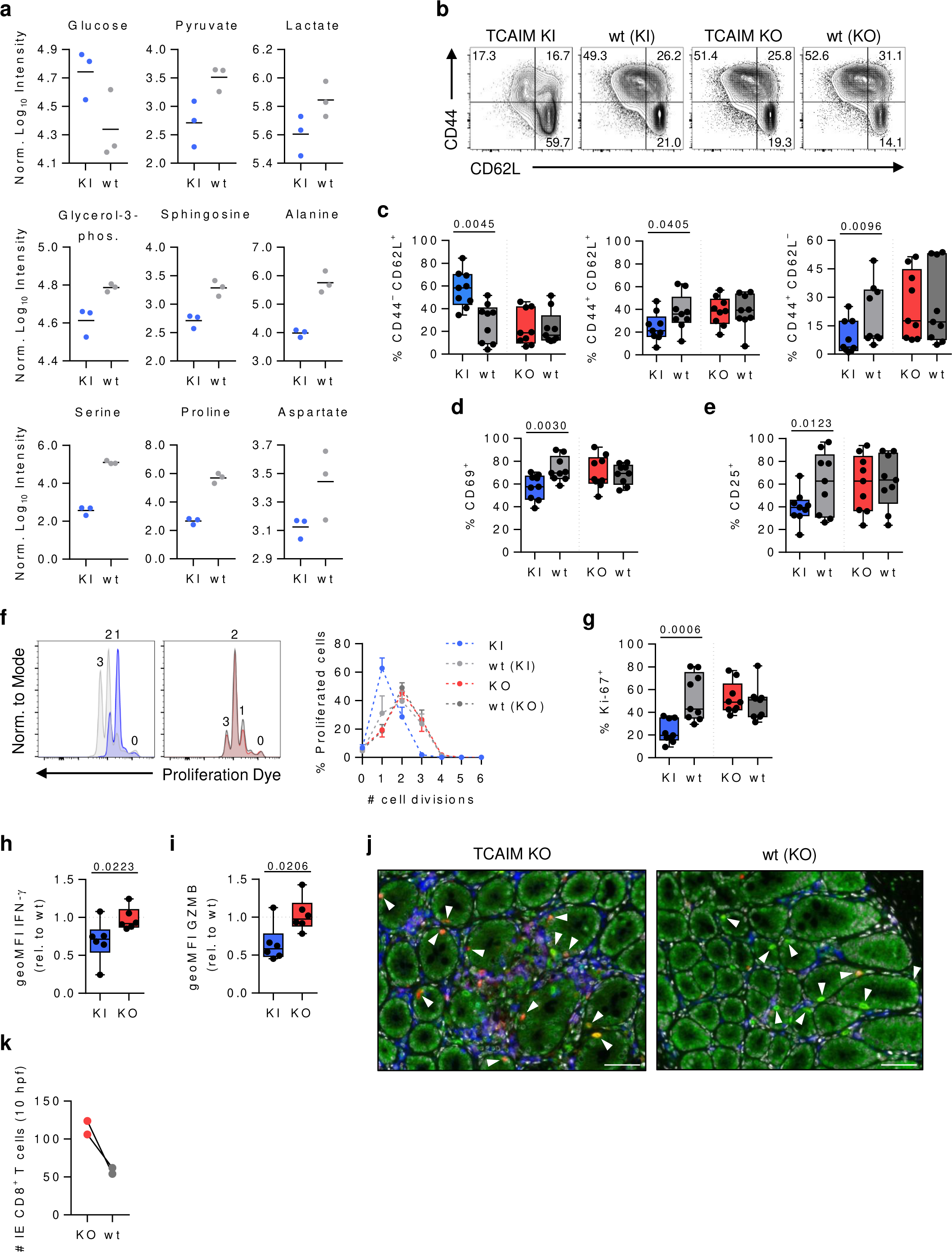
TCAIM KI alters the CD8^+^ T cell metabolic program and inhibits their proliferation and effector differentiation. (**a**) Non-targeted metabolite analysis of 60 h polyclonally activated TCAIM KI or wt CD8^+^ T cells (n = 3). Representative contour plots (**b**) and quantification of CD62L/ CD44 (**c**), CD69 (**d**) and CD25 (**e**) protein surface expression of 72 h activated TCAIM KI, KO and respective wt CD8⁺ T cells (n = 9). Cells were polyclonally activated with either 5 µg/ ml αCD3 and 2 µg/ ml αCD28 or 2 µg/ ml αCD3 and 1 µg/ ml αCD28. (**f**) Representative histogram overlays and quantification of in vitro proliferation of 48 h polyclonally activated TCAIM KI or wt CD8^+^ T cells co-cultured with B cells (n = 6). (**g**) Ki-67 protein expression of 48 h polyclonally activated TCAIM KI, KO and respective wt CD8⁺ T cells (n = 6). Intracellular IFN-γ (h) and GZMB (i) protein expression levels relative to wt controls of 48 h polyclonally activated TCAIM KI or KO CD8⁺ T cells co-cultured with B cells (n = 6). (j) Representative colon sections stained for CD3 (green), CD4 (blue) and CD8 (red) collected from T cell specific TCAIM KO mice or wt littermates. White arrowheads indicate epithelia infiltrating CD3^+^/ CD8^+^ T cells. Scale bars: 25 µm. (**k**) Absolute number of intraepithelial (IE) CD3^+^/ CD8^+^ T cells per 10 high power fields (hpf) in immunohistochemically stained colon sections (n = 2). Cells had been stimulated with 5 µg/ ml plate-bound αCD3 and 2 µg/ ml soluble αCD28 in a density of 6 x 10^5^ cells/ 200 µl culture medium (a,f-i). Quantitative data represents means (a), boxplots with whiskers from min to max (c-e,g-i), means ± SEM (f) or scatter dot plots (k). Data were analyzed by one-tailed, paired Student’s t-test (c-e), one-tailed, unpaired Student’s t-test (g-h) or one-tailed Mann-Whitney test (i). geoMFI, geometric mean fluorescence intensity.

Taken together, the non-targeted metabolite analysis revealed an altered metabolic program in activated TCAIM KI CD8^+^ T cells with reduced metabolite levels of glycolytic end products and biosynthesis pathways when compared to wt CD8^+^ T cells.

As a high glycolytic flux is essential for effector T cell differentiation, we next investigated whether TCAIM KI interferes with CD8^+^ T cell activation and differentiation. Again, naive TCAIM KI and wt CD8^+^ T cells were polyclonally activated and changes in CD62L and CD44 surface expression were measured by flow cytometry to discriminate between naïve (CD62L^+^ / CD44^−^), central memory (CD62L^+^ / CD44^+^) and effector/ effector memory (CD62L^−^ / CD44^+^) T cells. We additionally included CD8^+^ T cells isolated from mice with a conditional T cell specific knock-out (KO) of exon 4 in the Tcaim gene, which results in a loss of protein function, as well as strain specific Cd4-Cre wt controls to the analysis.

After 72 h of activation the majority of TCAIM KI CD8^+^ T cells maintained a naïve phenotype expressing CD62L only. Furthermore, the few activated TCAIM KI CD8^+^ T cells rather obtained a central memory phenotype expressing both CD62L and CD44. In contrast, TCAIM KO or wt CD8^+^ T cells were predominantly activated and differentiated into either effector/ effector memory or central memory cells (Fig. 1b,c).

We also analyzed expression of the two activation markers CD69 and CD25 indicative of the tissue homing and proliferative potential, respectively. Indeed, TCAIM KI also prevented upregulation of CD69 (Fig. 1d) and CD25 (Fig. 1e) expression in comparison to wt and TCAIM KO CD8^+^ T cells. However, CD69 and CD25 expression was not completely inhibited in TCAIM KI CD8^+^ T cells indicating residual T cell activation capacity. Nevertheless, this residual activation was not sufficient to acquire full proliferative potential (Fig. 1f,g) nor effector function (Fig. 1h,i).

Since TCAIM KI impairs CD8^+^ T cell activation and restricts differentiation, we expected an increased CD8^+^ T cell activation, proliferation and especially effector/ effector memory differentiation and function due to TCAIM KO. Yet, no significant differences between TCAIM KO and wt CD8^+^ T cells upon in vitro activation could be observed (Fig. 1b-i). However, ex vivo characterization of TCAIM KO CD8^+^ T cells by flow cytometry revealed a tendency towards increased spontaneous activation and effector/ effector memory differentiation of TCAIM KO CD8^+^ T cells in the colon (Extended Data Fig. 2a,b). Additionally, promoted a homozygous (hocre) TCAIM KO an age-dependent accumulation of effector/ effector memory CD8^+^ T cells in the colon, which could not be observed for heterozygous (hecre) or wild type (wtcre) TCAIM KO mice (Extended Data Fig. 2c). Furthermore, TCAIM KO mice showed an increased intra-epithelial infiltration rate of CD8^+^ T cells in the colon (Fig. 1j,k). We did not detect differences in frequencies and absolute numbers of total D8^+^ T cells in lymphatic or non-lymphatic tissues, e.g. colon, from TCAIM KO mice (Extended Data Fig. 2d,e).

**Fig. 2.**
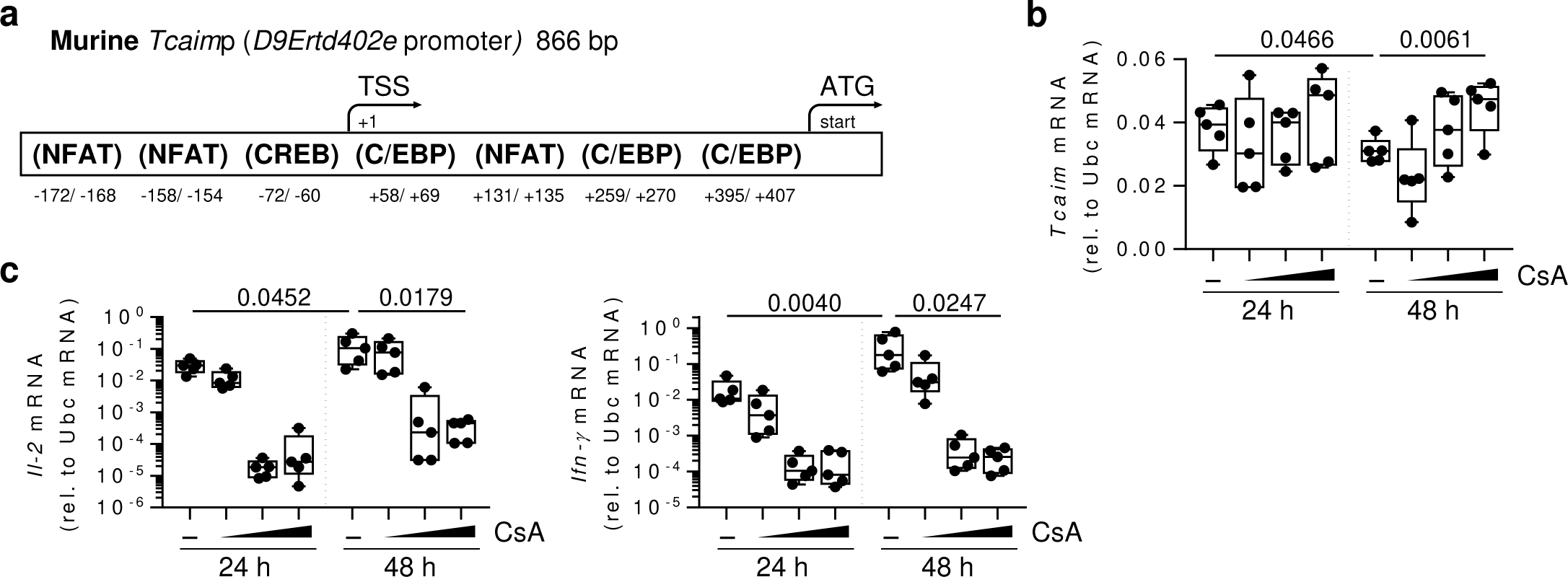
NFAT signaling regulates Tcaim gene expression. (**a**) Schematic representation of the murine D9Ertd402e promoter (Tcaimp) with predicted transcription factors binding sites displayed in brackets. Distance down- or upstream form the transcription start site (TSS) is indicated with - or +, respectively. Relative Tcaim (**b**) or Il-2 and Ifn-*γ* (**c**) mRNA expression analyzed by real-time qRT-PCR of 24 - 48 h polyclonally activated C57BL/6 CD8^+^ T cells upon treatment with the calcineurin inhibitor Cyclosporine A (CsA) or DMSO control (n = 5). Cells had been stimulated with 5 µg/ ml plate-bound αCD3 and 2 µg/ ml soluble αCD28 at a density of 6 x 10^5^ cells/ 200 µl culture medium. Gene expression was normalized to Ubc expression. Quantitative data represents boxplots with whiskers from min to max. Data were analyzed by one-tailed, unpaired Mann-Whitney test (Ifn-*γ* 24 vs. 48 h DMSO) or one-tailed, unpaired Student’s t-test (others).

Collectively, the data indicate that TCAIM KI leads to an altered metabolic program with reduced biosynthesis processes and suppressed CD8^+^ T cell activation and especially effector/ effector memory differentiation. In contrast, a TCAIM KO seems to be advantageous for CD8^+^ T cell activation, proliferation and intra-tissue accumulation of effector/ effector memory CD8^+^ T cells.

### Tcaim gene expression is regulated by NFAT

Knowing that TCAIM interferes with effector T cell differentiation and that its expression is downregulated upon activation^15^, we investigated the transcriptional control of its promoter. The promotor region of the murine Tcaim gene has predicted binding sites for nuclear factor of activated T cells (NFAT), CCAAT/enhancer-binding-proteine (C/EBP) and cAMP response element-binding protein (CREB) (Fig. 2a). Especially, NFAT function is essential for full T cell activation and acquisition of effector functions^17^. NFAT is activated by dephosphorylation through calcineurin^18^. Thus, we hypothesized NFAT as a negative regulator for Tcaim gene expression. To test this, we added the calcineurin inhibitor cyclosporine A (CsA) at increasing doses to stimulation cultures of naïve CD8^+^ T cells isolated from C57BL6/N mice.

### Tcaim mRNA expression was measured by real-time qPCR

DMSO control-treated CD8^+^ T cells showed a significant downregulation of Tcaim mRNA expression over time while CsA treatment resulted in a dose-dependent inhibition of Tcaim mRNA downregulation at 48 h of activation (Fig. 2b). Effective NFAT inhibition through CsA additionally was controlled by measuring mRNA expression of Il-2 and Ifn-*γ* mRNA as two main target genes of NFAT. Expression of both was significantly reduced with increasing doses of CsA, while untreated cells showed an increase in mRNA expression levels upon activation (Fig. 2c).

Taken together, NFAT indeed acts as a negative regulator for Tcaim gene expression.

### Metabolic modeling predicts forward directed glycolytic TCA cycle activity in general and reduced metabolic flux in TCAIM KI CD8^+^ T cells in particular

To further dissect the consequences of TCAIM KI for activation-induced upregulation of anabolic processes in CD8^+^ T cells, we analyzed the gene expression profile of naïve and polyclonally activated CD8^+^ T cells 24 to 48 h after stimulation from TCAIM KI, KO or wt mice by RNA-seq.

A targeted gene expression analysis of selected genes encoding for metabolically relevant transcription factors and enzymes of the glycolysis, tricarboxylic acid (TCA) cycle and glutaminolysis pathway revealed mainly a diminished upregulation of genes encoding for enzymes or enzyme subunits of the TCA cycle or glutaminolysis in both 24 and 48 h activated TCAIM KI compared to wt CD8^+^ T cells (Fig. 3a genes in bold).

**Fig. 3.**
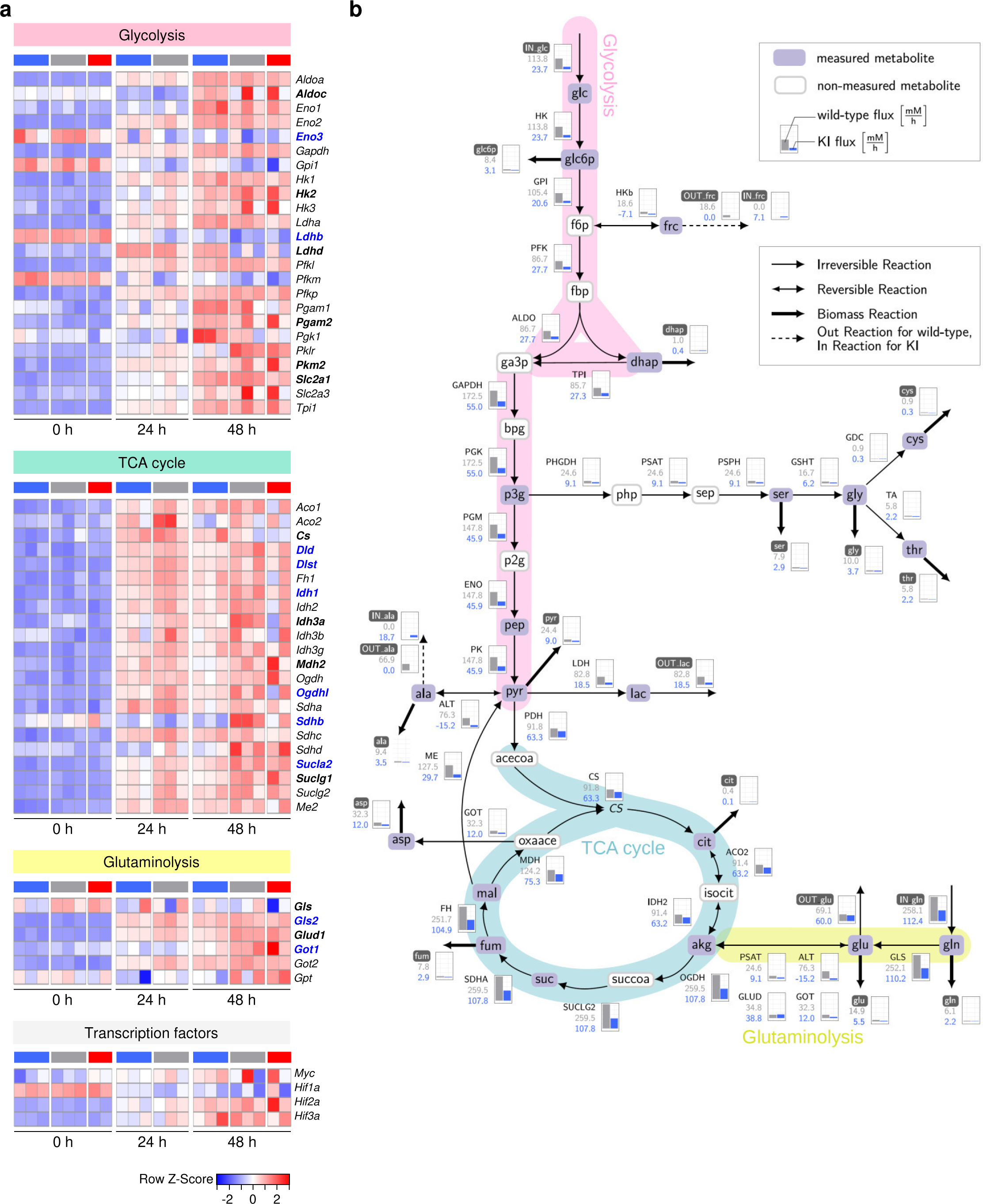
Reduced enzyme gene expression and metabolic flux rates in TCAIM KI CD8^+^ T cells. (**a**) Heatmap displaying normalized expression values (z-score) of selected genes encoding for glycolysis, tricarboxylic acid cycle and glutaminolysis controlling enzymes or transcription factors in naïve and 24 to 48 h polyclonally activated TCAIM KI (blue), KO (red) or wt (grey) CD8^+^ T cells (KI and wt: n = 3; KO: n = 2). Highlighted genes indicate those being differentially expressed between 48 h activated TCAIM KI and wt CD8^+^ T cells. P-values have been adjusted to either 0.05 (bold blue) or 0.3 (bold black). Expression of protein-coding genes was determined by normalization and variance-stabilized transformation of RNA-seq raw count data. (**b**) Mathematical metabolic flux modeling for 60 h polyclonally activated TCAIM KI or wt CD8^+^ T cells. Via mass spectrometry, measured metabolites are highlighted in purple boxes while non-measured metabolites are framed in grey. The flux distributions in mM / h according to the calculated enzyme activity of TCAIM KI and wt CD8^+^ T cells are displayed as bar graphs for each enzyme, respectively (wt: grey bar; KI: blue bar). Cells had been stimulated with 5 µg/ ml plate-bound αCD3 and 2 µg/ ml soluble αCD28 at a density of 6 x 10^5^ cells/ 200 µl culture medium.

The two main transcription factors for initiating and sustaining aerobic glycolysis and thus, activation-dependent metabolic adaptations of T cells, are MYC and HIF1α. Gene expression of both were upregulated in TCAIM KI CD8^+^ T cells and showed levels comparable to that of TCAIM KO and wt CD8^+^ T cells (Fig. 3a and Extended Data Fig. 3a). In accordance to upregulation to that, gene expression of glycolytic enzymes was upregulated in TCAIM KI, KO and wt CD8^+^ T cells 48 h after activation to a similar extend. This was unexpected, as we assumed a reduced glycolytic activity represented by a reduced glycolysis gene expression in TCAIM KI CD8^+^ T cells according to our previous observations of diminished glycolysis end product metabolite levels and abrogated effector differentiation. In contrast, there were genes with even significantly increased expression in TCAIM KI compared to wt CD8^+^ T cells either due to reduced down- or increased upregulation (e.g. Ldhd, Pgam2 or Ldhb, Pkm, Slc2a1, respectively). Only, Aldoc, Eno3 and the HIF1α target gene Hk2 were less upregulated in activated TCAIM KI CD8^+^ T cells with Eno3 being the only significant one (Fig. 3a). Differences in metabolic enzyme gene expression between activated TCAIM KO and wt CD8^+^ T cells could not be observed (Fig. 3a).

Thus, in accordance with the metabolite analysis, we observed reduced upregulation of TCA cycle and glutaminolysis but not for the majority of glycolysis controlling genes suggesting at least reduced metabolic rates in mitochondria of activated TCAIM KI CD8^+^ T cells.

The pleiotropic changes in the gene expression patterns cannot be directly linked to changes in metabolic fluxes, as the intermediate steps of protein translation and regulation of enzyme activity were not considered. We therefore mathematically constructed a model, which incorporated the available metabolomics data on TCAIM KI and wt CD8^+^ T cells to estimate their metabolic fluxes, with the TCAIM KI model being a differentially regulated variant of the wt model. The models shall foster the understanding of the general distribution of metabolic activity in activated CD8^+^ T cells, but also, the specific effects of TCAIM affecting this well-balanced process.

The calculated metabolic model of polyclonally activated wt CD8^+^ T cells revealed a strong support of the TCA cycle activity by increased glycolysis but also glutaminolysis rates (Fig. 3b grey bar charts of pink and yellow shaded paths). The TCA cycle flux was forward directed with no evidence of reversed citrate generation from glutamine to support biomass production. However, to ensure continuous forward TCA cycle flux, wt CD8^+^ T cells had to convert malate to pyruvate by malic enzyme (ME) to provide enough pyruvate and acetyl-CoA for increased citrate synthase (CS) activity (Fig. 3b grey ME bar chart). Similar to wt CD8^+^ T cells, activated TCAIM KI CD8^+^ T cells also showed an exclusively forward directed TCA cycle flux, which was driven by glycolysis and glutaminolysis (Fig. 3b blue bar charts of pink and yellow shaded paths). However, overall flux rates were strongly diminished in TCAIM KI CD8^+^ T cells (Fig. 3b blue bar charts). Also, differing to wt CD8^+^ T cells, glycolysis derived pyruvate was sufficient to meet their calculated CS activity and thus, showed no malate to pyruvate conversion.

In summary, the mathematical flux model suggests a continuous glycolytic TCA cycle flux in activated wt CD8^+^ T cells which was supported by increased glutaminolysis and malate to pyruvate conversion. In contrast, for activated TCAIM KI CD8^+^ T cells an overall reduced metabolic flux was proposed. However, except for the step of malate to pyruvate conversion directions of the major fluxes were not altered in comparison to wt CD8^+^ T cells.

### Decreased metabolic flux rates of activated TCAIM KI CD8^+^ T cells align with a reduced enzyme gene expression, ΔΨm and HIF1α protein expression

To validate the findings of the mathematically evaluated metabolic fluxes of TCAIM KI and wt CD8^+^ T cells, we compared gene expression of differentially expressed enzymes between polyclonally activated TCAIM KI and wt CD8^+^ T cells by RNA-seq (Extended Data Fig. 3b) with a regularization parameter introduced to calculate the deviating flux rates of TCAIM KI from that of wt CD8^+^ T cells. The regularization parameter describes the alteration of flux through each modelled enzyme summarizing the effects of different concentrations of enzyme substrates and products but also of co-factor availability. A negative regularization parameter value indicates a reduced activity of a particular enzyme and should align with a negative fold change in gene expression which indicates either a downregulation or reduced upregulation of this enzyme in TCAIM KI CD8^+^ T cells equivalent to a reduced enzyme activity. We thereby assume that mRNA levels correlate, at least to some extent, to enzyme activity.

As can be seen in Fig. 4a most of the enzyme regularization parameters aligned with the negative fold changes of the respective enzyme coding genes (Fig. 4a lower left corner). For the alanine transaminase (ALT) enzyme a regularization parameter for the backward flux rate (kb) was calculated, which is equivalent to a downregulated forward flux rate (kf). Thus, also the ALT regularization parameter agreed with a reduced Gpt2 (encoding for ALT) expression in TCAIM KI CD8^+^ T cells. The mismatches of the other calculated regulation parameters with their respective enzyme gene expressions in the upper left and lower right corners might be explained by missing information on either substrate and/ or product (GAPDH, SUCLG2 or PGM) as well as co-factor concentrations (MDH, PDH, GLUD, IDH2, LDH and PK). Especially the lack of knowledge about NAD^+^/ NADH^+^+H^+^ ratios in wt and TCAIM KI CD8^+^ T cells complicates a reliable regularization parameter calculation. However, assuming an increased NAD^+^/ NADH^+^+H^+^ ratio in TCAIM KI CD8^+^ T cells due to a decreased TCA cycle activity would decrease the regularization parameter of MDH, PDH, GLUD and IDH2 and increase the regularization parameter of LDH and, thereby, increase the chance of alignment with the fold changes in enzyme gene expression. Additionally, it should be kept in mind that enzyme gene expression is not perfectly ideal for metabolic flux modelling confirmation as many enzymes are regulated at a post-transcriptional level and may have other regulatory or structural functions such as GAPDH regulating IFN-Ȗ protein expression^2^.

**Fig. 4.**
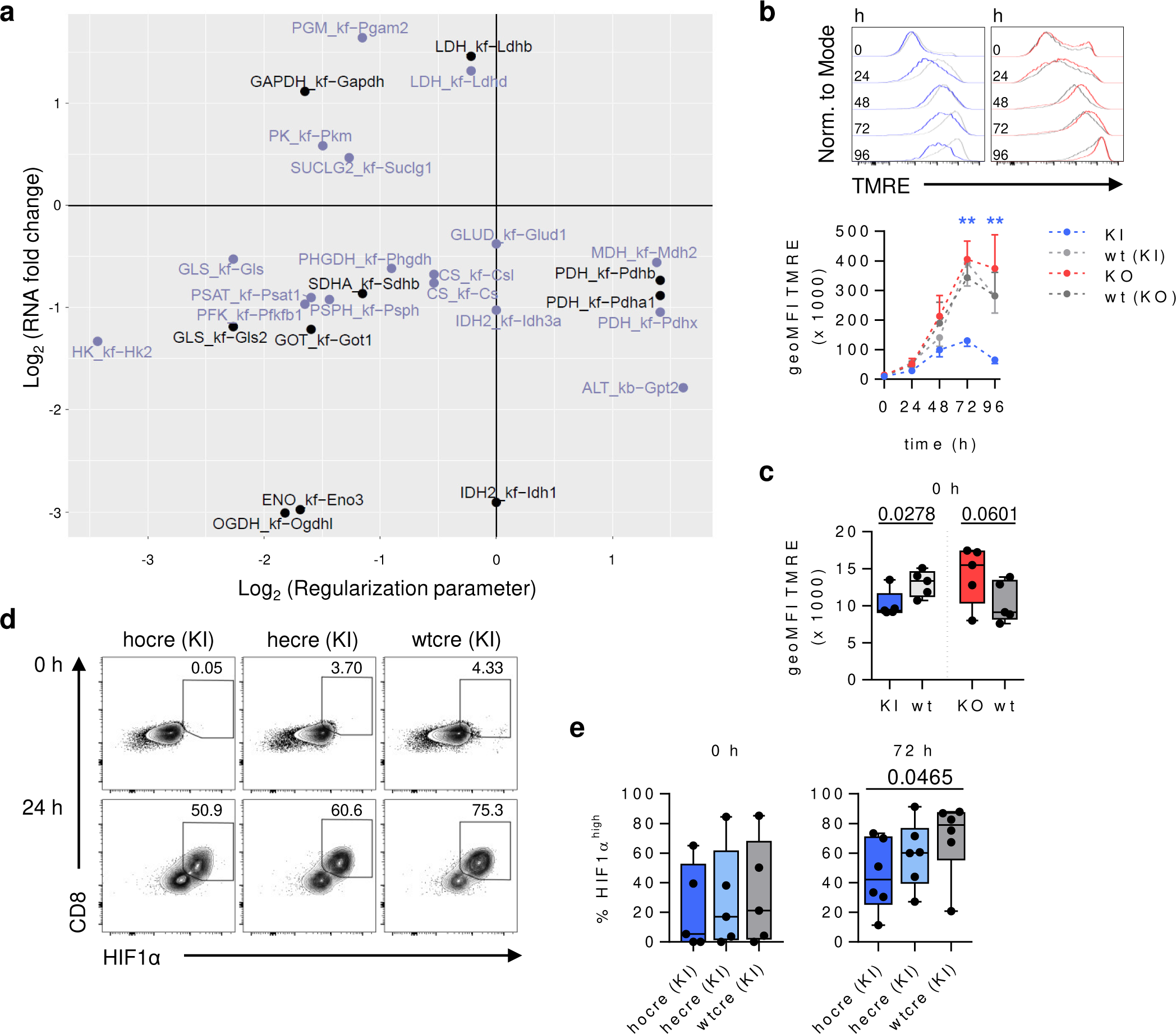
Reduced ΔΨm and HIF1α protein expression impairs metabolic activity in TCAIM KI CD8^+^ T cells. (**a**) Comparison of differentially expressed genes of 48 h polyclonally activated TCAIM KI or wt CD8^+^ T cells from RNA-seq analysis (log_2_ fold change) to regularization parameters introduced to describe the deviating TCAIM KI from the wt metabolic fluxes (log_2_). Only differentially expressed genes with adjusted p-values of ≤ 0.05 (black) or ≤ 0.3 (purple) are included. Enzymatic fluxes are described with either forward (kf) or backward (kb) flux rates. (**b**) Representative histogram overlays and quantification of TMRE staining for ΔΨm of naïve and polyclonally activated TCAIM KI, KO and respective wt CD8^+^ T cells co-cultured with B cells (n = 5). Significant differences between TCAIM KI or KO and their wt controls are indicated with blue or red asterisks, respectively. (**c**) TMRE staining for baseline ΔΨm of naïve TCAIM KI, KO and respective wt CD8^+^ T cells co-cultured with B cells (n = 5). Representative contour plots (**d**) and quantification (**e**) of HIF1α expression of naïve and 72 h polyclonally activated homozygous (hocre), heterozygous (hecre) TCAIM KI or wild type (wtcre) CD8⁺ T cells (n = 6). Cells had been stimulated with 5 µg/ ml plate-bound αCD3 and 2 µg/ ml soluble αCD28 at a density of 6 x 10^5^ cells/ 200 µl culture medium (a). Quantitative flow cytometry data represents means ± SEM (b) or boxplots with whiskers from min to max (c,e). Data were analyzed according to Gaussian distribution of the compared sample sets and for each genotype with its respective wt control using (b) Kruskal-Wallis test with Dunn’s multiple comparisons test (KI vs. wt) or one-way ANOVA (KO vs. wt) and subsequent post hoc one-tailed Mann-Whitney test (KI vs. wt 0 h) or one-tailed, unpaired Student’s t-test (others), (c) using one-tailed Mann-Whitney test (KI vs wt) or one-tailed, unpaired Student’s t-test (KO vs wt) or (e) Kruskal-Wallis test with Dunn’s multiple comparisons test and subsequent post hoc one-tailed, unpaired Mann-Whitney test. * p < 0.05, ** p < 0.01. geoMFI, geometric mean fluorescence intensity.

The integrity of ΔΨm is closely linked to the TCA cycle and OXPHOS activity. NADH+H^+^ generated by the TCA cycle is used as electron donor by the respiratory complexes^19^. In consequence of the electron transport, a proton gradient is built up, which is reflected by the ΔΨm and used for ATP synthesis by OXPHOS^20^. As we observed a reduced TCA cycle activity in the metabolic flux model of TCAIM KI CD8^+^ T cells, we hypothesized that they also have a lower ΔΨm. We performed flow cytometry analysis with naïve and polyclonally activated TCAIM KI, KO and wt CD8^+^ T cells and measured the ΔΨm using the fluorescent dye TMRE.

All CD8^+^ T cells responded to the activation stimulus with an increase of the ΔΨm reaching a peak after 72 h, but as expected, the increase was significantly less in TCAIM KI CD8^+^ T cells (Fig. 4b). Additionally, the baseline ΔΨm was also significantly lower in TCAIM KI CD8^+^ T cells compared to the respective wt control while it tended to be higher in TCAIM KO CD8^+^ T cells (Fig. 4c).

A prolonged high glycolytic flux in activated T cells is controlled by the transcription factor HIF1α^4^. To ensure rapid adaptation to changes in oxygen availability or immune stimulation HIF1α is constantly expressed. Under homeostatic conditions in non-activated T cells the protein is immediately degraded, but upon T cell activation HIF1α mRNA and protein expression becomes upregulated and HIF1α protein degradation is inhibited^21^. The latter is especially linked to increased levels of the TCA cycle metabolites succinate^22^ and fumarate^23^ but also to increased mROS level^21^.

As mentioned before, Hif1a mRNA expression was upregulated upon activation in both TCAIM KI and wt CD8^+^ T cells. However, as HIF1α is primarily regulated at the post-transcriptional level and the mathematical metabolic modelling of TCAIM KI CD8^+^

T cells predicted a reduced TCA cycle activity correlating with reduced levels of TCA cycle derived succinate and fumarate, we hypothesized that HIF1α protein expression was reduced upon activation of TCAIM KI CD8^+^ T cells. Thus, we measured HIF1α protein expression by flow cytometry. Indeed, TCAIM KI CD8^+^ T cells showed significantly decreased HIF1α protein levels upon activation compared to wt CD8^+^ T cells in a gene dose depend manner (Fig. 4d,e).

Collectively, the crosscheck of the mathematical model with the RNA-seq data revealed predominantly the agreement of decreased enzyme activities with reduced enzyme gene expression in TCAIM KI CD8^+^ T cells. Moreover, TCAIM KI CD8^+^ T cells seem to fail to increase their ΔΨm as well as their HIF1α protein expression upon activation further confirming our assumption of impaired metabolic activity and thus, effector differentiation due to TCAIM KI.

### TCAIM inhibits cholesterol biosynthesis in activated CD8^+^ T cells

As we observed a regulatory role of TCAIM for metabolic adaptations upon T cell activation, we investigated whether there was a mechanistic explanation evident from our RNA-seq data. In a non-supervised hierarchical clustering analysis naïve CD8^+^ T cells clearly separated from all activated T cells. TCAIM KI or KO had no further effect on the subclustering of naïve CD8^+^ T cells. Interestingly, the expression profile of 48 h activated TCAIM KI CD8^+^ T cells remained more similar to that of 24 h activated TCAIM KI and wt CD8^+^ T cells, whereas 48 h activated TCAIM KO and wt CD8^+^ T cells acquired a distinct expression profile (Fig. 5a).

**Fig. 5.**
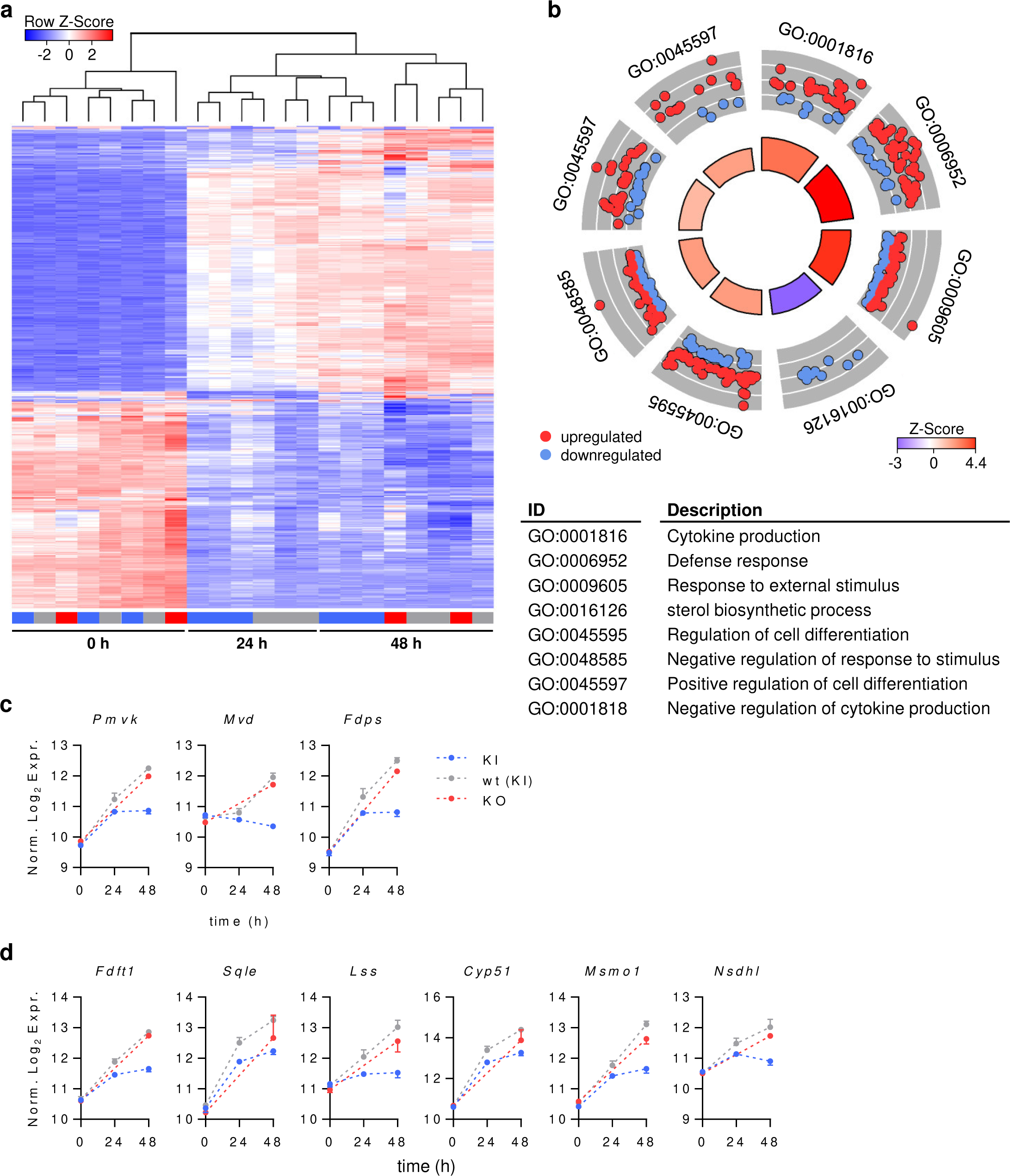
TCAIM inhibits activation-induced expression of cholesterol biosynthesis genes. (**a**) Non-supervised hierarchical clustering analysis of 1000 genes with the highest variances across naïve and polyclonally activated TCAIM KI (blue), KO (red) or wt (grey) CD8^+^ T cells (KI and wt: n = 3; KO: n = 2). Normalized expression values (z-scores) are depicted. Expression of protein-coding genes was determined by normalization and variance-stabilized transformation of RNA-seq raw count data. (**b**) DAVID functional gene ontology (GO) enrichment analysis within the sets of differentially expressed genes between 48 h polyclonally activated TCAIM KI or wt CD8^+^ T cells (n = 3). GO circle plot displaying representative GO terms selected out of the top 25 GO biological processes. The outer ring shows the log_2_ fold change (FC) of the assigned genes from higher to lower (outer to inner layer) for each term. Red circles display genes being upregulated and blue ones being downregulated. The inner ring shows the z-Score [(#genes_up_ - #genes_down_) / (#genes_up_ + #genes_down_)] corresponding to the blue red gradient. The width of the inner ring bars indicate the significance of each GO term in relation to each other – the wider the bar, the smaller the p value (log_10_-adjusted p-value < 0.05). mRNA expression of enzymes controlling the mevalonate pathway (**c**) and cholesterol biosynthesis (**d**) of naïve and polyclonally activated TCAIM KI, KO or wt CD8^+^ T cells (KI and wt: n = 3; KO: n = 2). RNA-seq raw count data of protein-coding genes were normalized and log_2_-transformed after adding a pseudocount to each value. Cells had been stimulated with 5 µg/ ml plate-bound αCD3 and 2 µg/ ml soluble αCD28 at a density of 6 x 10^5^ cells/ 200 µl culture medium. Quantitative data represent means ± SEM (c-d).

To understand the functional differences associated with the altered expression profile in activated TCAIM KI CD8^+^ T cells, we performed a gene ontology (GO) enrichment analysis in which genes were differentially expressed between 48 h activated TCAIM KI and wt CD8^+^

T cells (Fig. 5b). The majority of annotations found among the GO terms describing biological processes were related to immunologic processes including cytokine production, responses to stimuli or cell differentiation. Genes that repeatedly contributed to that annotations comprised e.g. Klf2, Il-1r2, Traf3ip1, Ifng-*γ*, Klrk1, IL-23a, Cd80 or Il-12rb (Extended Data Tab. 1) with the latter five as markers for T cell activation and effector function being downregulated, while the others as markers for T cell quiescence and inhibition being upregulated in TCAIM KI CD8^+^ T cells. This is again consistent with the impaired T cell activation and effector function due to TCAIM KI.

Remarkably, apart from the regulation of genes controlling immunologic processes we detected a strong downregulation of genes involved in cholesterol biosynthetic processes. Genes within this set were essential enzymes of the mevalonate pathway (Fig. 5c) and the cholesterol biosynthesis which branches off the mevalonate pathway (Fig. 5d). Indeed, TCAIM KI CD8^+^ T cells either failed to upregulate gene expression at all (e.g Mvd or Lss) or to further increase gene expression after 24 h of activation (Pmvk, Fdps, Fdft1, Sqle, Cyp51, Msmo1 and Nsdhl; Fig. 5c,d). This was also true for most of the remaining enzymes of the mevalonate and cholesterol biosynthesis pathways although their changes were not significant (Extended Data Fig. 4a,b).

Taken together, the RNA-seq analysis further confirmed an impaired T cell activation and acquisition of effector functions and additionally, revealed a deficiency in expression of cholesterol biosynthesis genes due to TCAIM KI.

### Cholesterol supplementation restores activation and proliferation of TCAIM KI CD8^+^T cells

As TCAIM KI CD8^+^ T cells showed a reduced cholesterol biosynthesis based on the RNA-seq data, we tested whether replenishment of free cholesterol could restore CD8^+^ T cell activation and especially effector differentiation. To investigate this, naïve CD8^+^ T cells with homozygous, heterozygous TCAIM KI or wt TCAIM gene expression (hocre, hecre or wtcre, respectively) were cultured in presence of increasing amounts of cholesterol for up to 72 h.

We then performed flow cytometry analysis and measured CD62L, CD44 and HIF1α protein expression as well as CD8^+^ T cell proliferation.

CD8^+^ T cell activation and differentiation clearly depended on the level of TCAIM overexpression. The majority of untreated hocre TCAIM KI CD8^+^ T cells again maintained a naïve phenotype expressing CD62L only, whereas untreated hecre TCAIM KI CD8^+^ T cells showed an increase in central memory and effector/ effector memory frequencies. Those, however, still were below that of untreated wtcre CD8^+^ T cells. Remarkably, this changed upon addition of cholesterol in a dose-dependent manner. We detected a significant increase in central memory cell frequencies upon treatment with the highest cholesterol dose for hocre TCAIM KI CD8^+^ T cells, which were then comparable to the levels of untreated wtcre CD8^+^

T cells. A similar tendency although not significant could also be observed for the effector/ effector memory cell pool of hocre as well as hecre TCAIM KI CD8^+^ T cells. Wtcre CD8^+^ T cells, however, did not respond to the cholesterol treatment and showed no further activation (Fig. 6a,b). The increased activation upon cholesterol treatment was accompanied by a likewise increased HIF1α protein expression and proliferation rate of both hocre and hecre TCAIM KI but not wt CD8^+^ T cells (Fig. 6c-e).

**Fig. 6.**
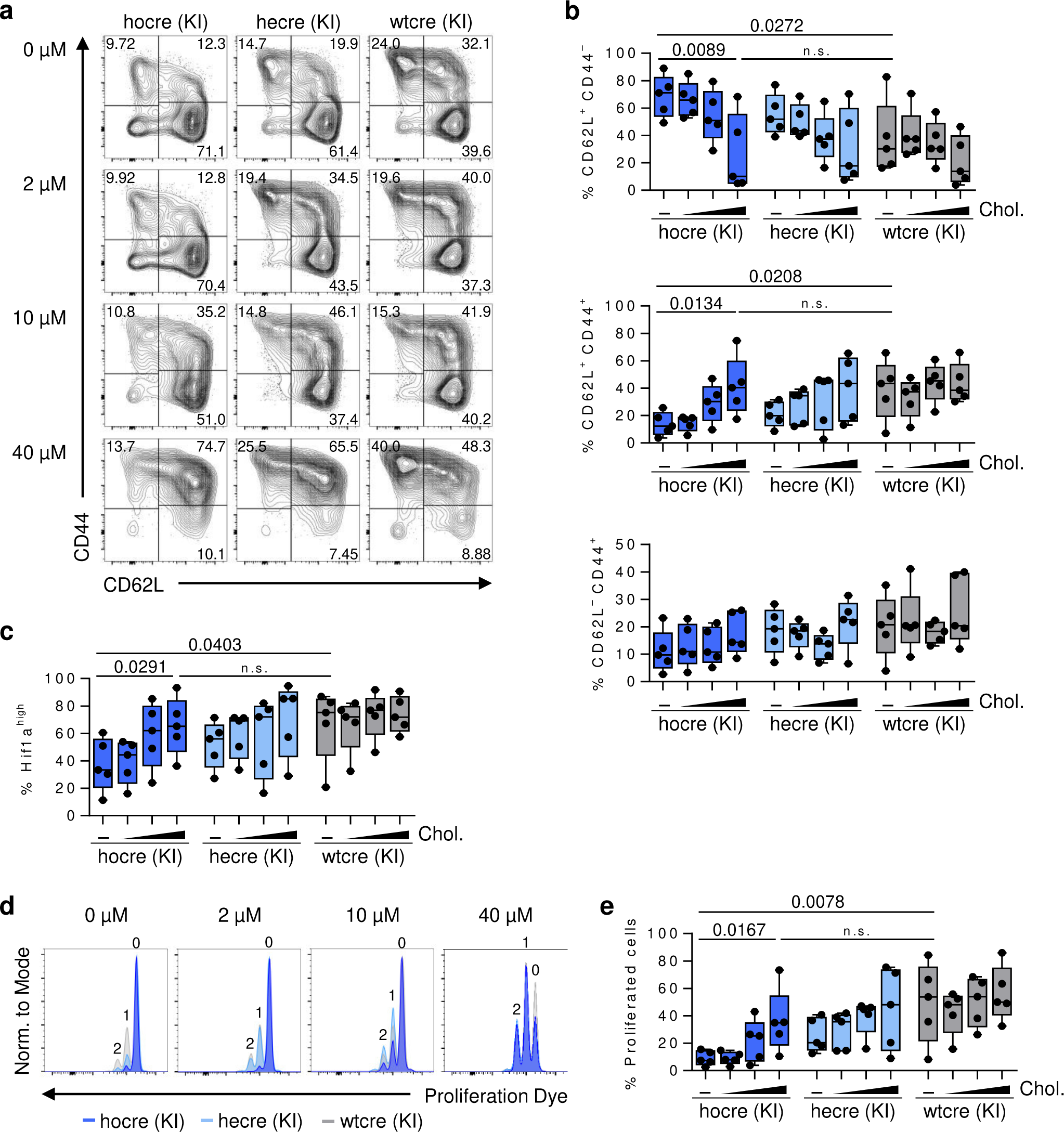
Exogenous cholesterol restores activation and proliferation of TCAIM KI CD8^+^ T cells. Representative contour plots (**a**) with quantification of CD62L / CD44 surface protein expression (**b**), intracellular HIF1α protein expression (**c**) and representative histogram overlays (**d**) with quantification of in vitro proliferation (**e**) of 72 h polyclonally activated homozygous (hocre), heterozygous (hecre) TCAIM KI or wild type (wtcre) CD8⁺ T cells (n = 5). Cells have been treated with increasing doses of Cholesterol or have been left untreated as indicated. Quantitative data represents boxplots with whiskers from min to max and were analyzed by one way ANOVA with post hoc one-tailed, unpaired Student’s t-test. n.s., not significant (p > 0.05).

In summary, the addition of exogenous cholesterol was sufficient to restore the activation, proliferation and partially differentiation deficiency of TCAIM KI CD8^+^ T cells. Thus, we conclude that TCAIM mediated defects in CD8^+^ T cell activation and differentiation were due to a decrease in cholesterol biosynthesis enzyme expression.

### TCAIM controls activation-induced MERC formation

Mitochondrial fission accompanied cristae remodeling was proposed as prerequisite for activation-induced metabolic adaptations and finally acquisition of effector function^13^. As TCAIM KI CD8^+^ T cells showed an altered metabolic program and impaired effector differentiation, we next investigated whether TCAIM regulates mitochondrial dynamics and cristae remodeling. Thus, we analyzed electron microscopy images of naïve (0h) and 24 - 48 h polyclonally activated TCAIM KI, KO and wt CD8^+^ T cells (Fig. 7a and Extended Data Fig. 5a).

**Fig. 7.**
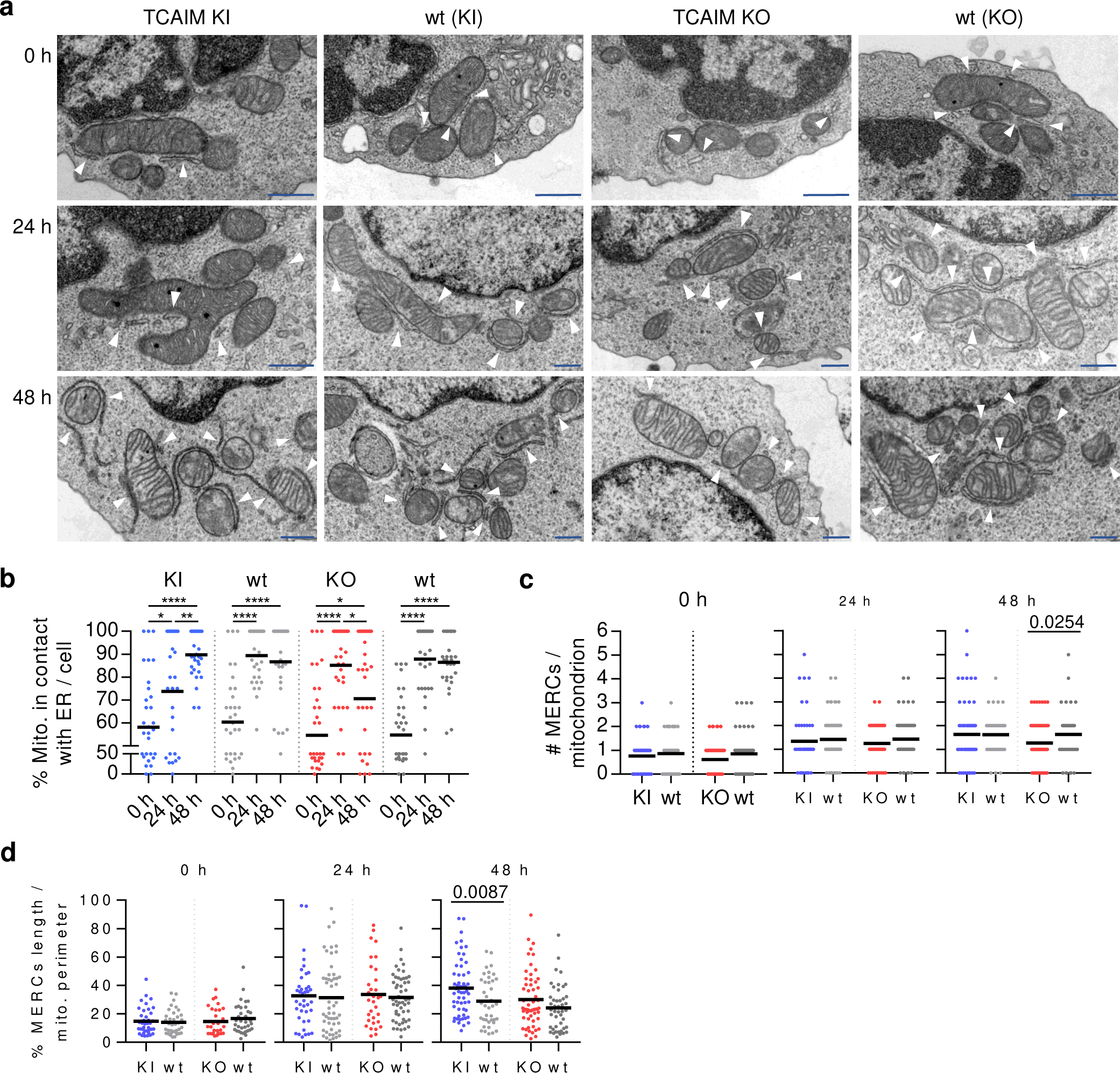
TCAIM KI delays MERC formation. (**a**) Zoomed regions of representative transmission electron microscopy images of naïve (upper row) and 24 to 48 h (middle and bottom row, respectively) polyclonally activated TCAIM KI, KO or respective wt CD8^+^ T cells. White arrowheads indicate mitochondria-ER interaction sites. Scale bars: 500 nm. Digitally magnification: 2.35 x. (**b-d**) Quantitative plots of transmission electron microscopy image analysis of naïve and polyclonally activated TCAIM KI, KO and respective wt CD8^+^ T cells. (**b**) Frequency of mitochondria being in contact with the ER per cell (n = 30). (**c**) Absolute number of mitochondria-ER contact sites (MERCs) per mitochondrion. (**d**) Proportional share of total MERCs length’s to mitochondrial perimeter per mitochondrion. For (c) and (d) a minimum of seven cells with a minimum of 30 mitochondria being in contact with the ER have been analyzed. If there were less than 30 mitochondria with ER contact sites the number of cells being analyzed was increased to a maximum of up to 9 cells.

The data revealed an increase in the average number of mitochondria per cell upon activation of wt but also TCAIM KI as well as TCAIM KO CD8^+^ T cells. This is in line with previous findings on human and murine CD8^+^ T cells, which showed that the number of mitochondria per cell increased upon activation^24, 25^. However, while the increase in number of mitochondria was fast but only transient for wt and TCAIM KO CD8^+^ T cells, TCAIM KI CD8^+^ T cells showed a delayed increase in mitochondrial number (Extended Data Fig. 5b). Against expectations, no structural differences in mitochondrial shape towards a more elongated morphology pointing towards a less anabolic phenotype could be observed for TCAIM KI CD8^+^ T cells. Also vice versa, TCAIM KO did not cause an increased fission rate in TCAIM KO CD8^+^ T cells (Extended Data Fig. 5c). However, in line with the increase in average mitochondrial number, we observed an activation-dependent transient increase in cristae formation for all TCAIM KI, KO and wt CD8^+^ T cells (Extended Data Fig. 5d). Interestingly, we detected disparities of MERCs between activated TCAIM KI, KO and wt CD8^+^ T cells (Fig. 7a). On average, 60 % of the mitochondria from a single naïve CD8^+^ T cell were in contact with the ER. This increased dramatically upon activation with up to 90 % of the mitochondria from wt and TCAIM KO CD8^+^ T cells engaging contact with the ER 24 h after stimulation (Fig. 7b). In contrast, mitochondria from TCAIM KI CD8^+^ T cells displayed a delayed MERC response with 70 % and 90 % of the mitochondria forming contacts at 24 and 48 h, respectively. Whereas, MERCs of wt and most strikingly TCAIM KO CD8^+^ T cells began to dissolve 48 h after stimulation as only 85 % and 70 % of the mitochondria, respectively, had MERCs left (Fig. 7b). Also, the number as well as the length of single ER tubule being in contact with a mitochondrion increased upon activation (Fig. 7c,d). Importantly, TCAIM KO CD8^+^ T cells had significantly less ER tubule contacts per mitochondrion (Fig. 7c) while the ER tubule contacts formed within TCAIM KI CD8^+^ T cells had a highly increased length and shape with some mitochondria being nearly completely covered by ER tubules (Fig. 7d) after 48 h of stimulation.

Collectively, activated TCAIM KI CD8^+^ T cells showed a delayed but strong increase in number of mitochondria and delayed formation of very long MERCs. No clear evidence for promotion of mitochondrial fusion was found due to TCAIM KI. Thus, the altered anabolic adaptation of activated TCAIM KI CD8^+^ T cells may not be caused by changed mitochondrial dynamics in general but by delayed and altered MERC formation.

### TCAIM interacts with proteins being enriched at MERC sites

The interaction between mitochondria and ER has been described to regulate mitochondrial metabolism and bioenergetics, mitochondrial fission, lipid metabolism, cellular Ca^2+^ signaling, autophagosome formation and apoptosis^26, 27^. To mediate these processes, mitochondria and ER membranes need to physically tether. This is accomplished by specific protein interactions, which are not yet fully identified for mammalian and especially immune cells. Protein pairs that were previously characterized to actively mediate mitochondria-ER tethering include mitochondrial MFN1 and MFN2 interacting with ER bound MFN2, VDAC2 with IP3R and Gpr75 or RMD3 with VAPB, respectively ^27^. Thus, to get a more detailed information on how TCAIM interferes with MERC formation we identified its interaction partners. Hence, we performed a anti-GFP co-immunoprecipitation with cell lysates from HEK293T cells transfected with either a plasmid encoding for a TCAIM-eGFP fusion protein or an eGFP protein with mitochondrial targeting sequence as a control. Subsequent mass spectrometry of the retained co-immunoprecipitates revealed only 14 TCAIM specific interaction partners (Fig. 8a). These include TCAIM itself suggesting formation of homomers to exert TCAIM specific functions. The other identified interaction partners were mostly proteins known to be located to mitochondria. More specifically, six of the TCAIM interaction partner, including TOM40, SAM50, VDAC2, MIC60, DJC11 and STML2, belong to the mitochondrial contact site and cristae organizing system (MICOS) protein complex and MICOS-associated proteins. MICOS and MICOS-associated proteins have been shown to regulate cristae assembly, but also, to participate in MERC formation, where they mediate the segregation of inner mitochondrial membrane components for appropriate cristae, protein and mtDNA distribution^28^. Notably, the previously characterized mitochondria-ER tethering proteins VDAC2 and RMD3 were among the identified TCAIM interaction partners.

**Fig. 8.**
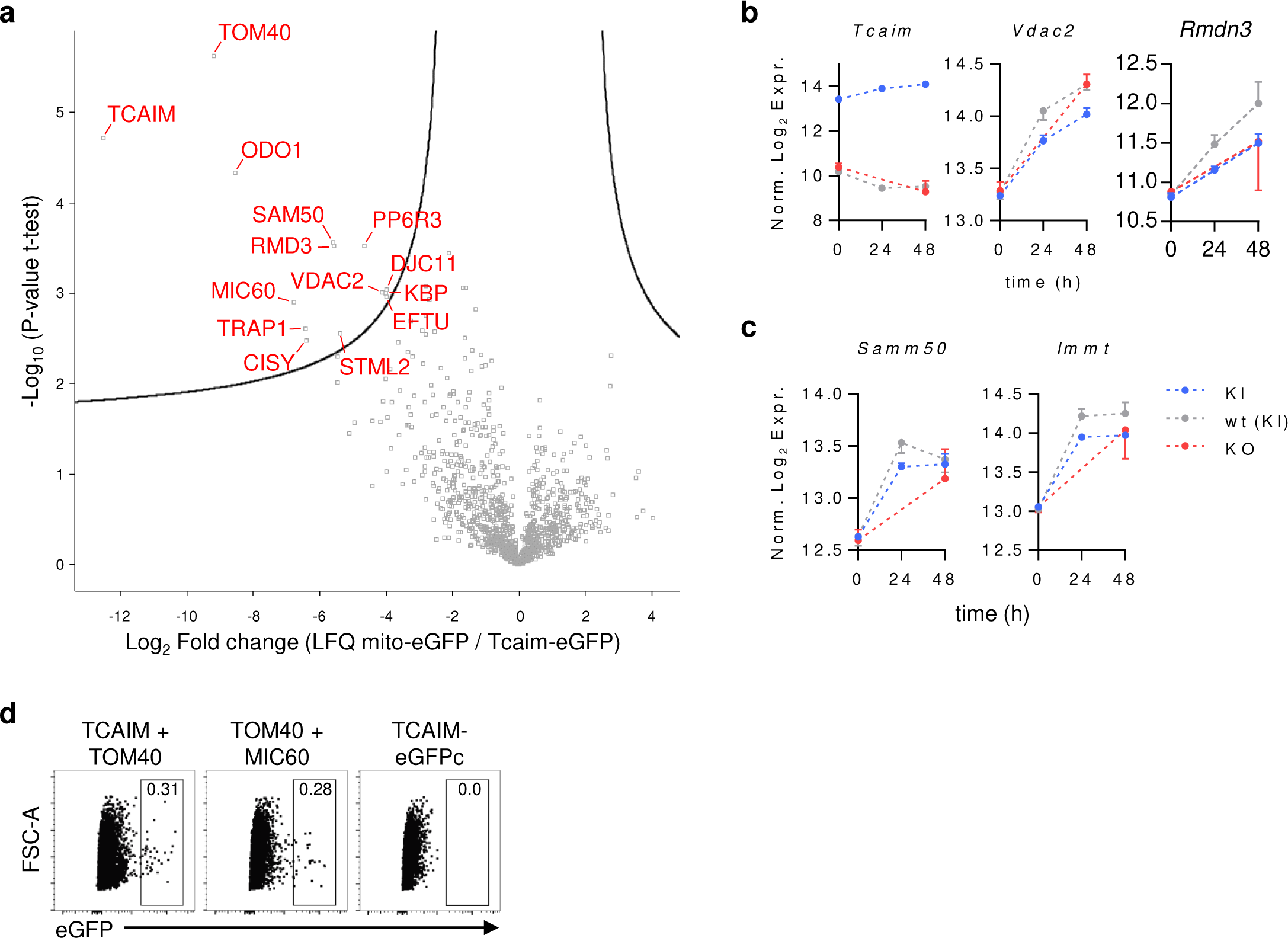
TCAIM interacts with MERC promoting proteins. (**a**) GFP co-immunoprecipitates of cell lysates from HEK293T cells transfected with pTCAIM-EGFP-N1 or mito-PAGFP control plasmid were analyzed for TCAIM interaction partner by LC-MS/MS. Volcano plot showing 14 proteins (FDR 0.05, S0 0.7) being significantly enriched only in pTCAIM-EGFP-N1 cell lysate fractions (n = 3). (**b-c**) mRNA expression of selected TCAIM interaction partners of naïve and polyclonally activated TCAIM KI, KO or wt CD8^+^ T cells (KI and wt: n = 3; KO: n = 2). RNA-seq raw count data of protein-coding genes were normalized and log_2_-transformed after adding a pseudocount to each value. Cells had been stimulated with 5 µg/ ml plate-bound αCD3 and 2 µg/ ml soluble αCD28 at a density of 6 x 10^5^ cells/ 200 µl culture medium. (**d**) Representative dot plots from one out of two BiFC assay experiments for validation of TCAIM interaction with TOM40 measured by flow cytometry. As indicated HEK293T cells were transfected with plasmids encoding for the split eGFP fusion proteins TCAIM-eGFPc + TOM40-eGFPn, TOM40-eGFPn + MIC60-eGFPc as positive control or TCAIM-eGFPc alone as negative control. Quantitative data represents means ± SEM (b-c).

Gene expression of both proteins (Rmdn3 encodes for RMD3) was upregulated upon activation in all TCAIM KI, KO and wt CD8^+^ T cells, whereas TCAIM gene expression was only downregulated in TCAIM KO and wt but not TCAIM KI CD8^+^ T cells (Fig. 8b). Despite VDAC2 and RMD3, also gene expression of SAM50 and MIC60, core-subunits of the MICOS protein complex, was upregulated in TCAIM KI, KO and wt CD8^+^ T cells (Fig. 8c).

Finally, bimolecular fluorescence complementation (BiFC) assay with subsequent flow cytometric analysis was used to confirm interaction of TCAIM with the MICOS subunit TOM40. The assay is based on the reconstitution of a fluorescent protein. HEK293T cells were transfected with two plasmids each encoding for the indicated protein of interest fused to either the N- or C-terminal part of eGFP. In case of protein interaction, the complementary eGFP fragments refold and start to fluoresce. Transfection of TCAIM fused to C-terminal part of eGFP alone was used as control for false positive signals like auto fluorescence of HEK293T cells while transfection of plasmids encoding for TOM40 and MIC60 which are known to interact with each other^29^ within the MICOS complex served as positive control. Indeed, eGFP positive cells were observed when complementary eGFP fragments coupled to TCAIM and TOM40 as well as TOM40 and MIC60 were transfected (Fig. 8d).

In conclusion, our results revealed that TCAIM interacts with proteins promoting MERC formation and function. Moreover, our results show that T cell activation enhances transcription of MERC forming proteins, whereas, down-regulation of TCAIM expression might circumvent inhibition of MERC formation.

## Discussion

T cells need to adapt their cellular metabolism for full effector cell differentiation upon activation and it has been reported that this relies on alterations in mitochondrial physiology including cristae resolution, fission and an increase in ΔΨm and mROS production^11, 13^.

However, the molecular signals driving mitochondrial changes and the communication between mitochondria and the ER thereby mediating changes in cellular metabolism remain less clear. We recently reported that the mitochondrial protein TCAIM is downregulated upon T cell activation and inhibits effector cell differentiation of conventional CD4^+^ T cells^15^. Our previous investigations also indicated that TCAIM interferes with activation-induced changes in mitochondrial morphology and function^15^. Thus, TCAIM seems to function as an important link between activation-induced mitochondrial alterations and changes in cellular metabolism driving full effector cell differentiation.

Therefore, we here utilized conditional TCAIM KI and KO mice as a model to investigate which molecular processes in the mitochondria control effector T cell differentiation: First, we could show that TCAIM KI causes also in CD8^+^ T cells comparable effects of impaired proliferation, effector/ effector memory differentiation and cytokine production as we previously reported for CD4^+^ T cells^15^. Second, our data on TCAIM KO CD8^+^ T cells point towards an advantage for effector/ effector memory differentiation in vivo. This further supports the regulatory role of TCAIM for activation and differentiation of T cells in general. Third, metabolomics and transcriptional data revealed that TCAIM KI inhibits activation-induced enforcement of TCA cycle activity and glutaminolysis, but more importantly, that TCAIM KI blocks upregulation of the mevalonate pathway and the cholesterol biosynthesis. Indeed, addition of cholesterol rescued T cell activation and proliferation of TCAIM KI CD8^+^ T cells. Finally, at the molecular level, we could show that TCAIM delays activation-induced interaction between mitochondria and the ER probably by binding to proteins known to promote MERC formation such as RMD3 and VDAC.

For a long time, many studies on activation-induced metabolic adaptations in T cells were primarily focusing on the glycolysis and TCA cycle axis as the central carbon processing pathways. However, enforcement of lipid synthesis following activation has been shown to be likewise important for T cell proliferation and acquisition of effector functions. Chen et al.^30^ already found in the 1970^th^, a strong upregulation of cholesterol biosynthesis in lymphocytes within the first 24 h upon activation and showed the importance for induction of DNA replication and cell proliferation. Later, in the 1990^th^, Chakrabarti et al.^31^ showed, that T cells were unable to proceed in cell cycle progression upon inhibition of cholesterol biosynthesis and instead remained in G1 phase. In addition to promoting T cell proliferation, Kidani et al.^32^ reported on the importance of cholesterol stores and biosynthesis for virus specific effector cell development, whereas Yang et al.^33^ demonstrated the need of increased membrane cholesterol during T cell activation for enhancement of TCR signaling and immunologic synapse formation.

Our study not only supports the importance of cholesterol for CD8^+^ T cell activation and proliferation, but also revealed a crucial role of activation-induced MERC formation which was inhibited by TCAIM KI. Notably, cholesterol biosynthesis enzymes are located in MERC structures^34^ further highlighting the role of mitochondria-ER communication for cholesterol biosynthesis. We could show that TCAIM binds to MERC promoting proteins such as VDAC and RMD3, which are crucial for MERC formation and function including lipid biosynthesis^35–38^.

Moreover, we showed that TCAIM interferes with increased HIF1α protein expression upon T cell activation, which could be restored upon addition of exogenous cholesterol.

Cholesterol treatment of hepatocytes has been shown to induce HIF1α protein accumulation mediated by inhibition of HIF1α protein degradation^39^. Furthermore, HIF1α protein expression as well as induction of CD44 is promoted by mammalian target of rapamycin complex 1 (mTORC1)^40, 41^. MTORC1 also promotes SREBP2 activation^42, 43^ and protein expression^32^ which is crucially controlling transcription of mevalonate pathway and cholesterol biosynthesis enzymes^44^. Indeed, we found reduced cholesterol biosynthesis enzyme gene expression as well as HIF1α and CD44 protein expression in activated TCAIM KI CD8^+^

T cells suggesting a deficiency in mTORC1 signaling through TCAIM mediated mechanisms. Although not explicitly shown for T cells, mTORC1 has been found to be located to mitochondria thereby regulating mitochondrial functions^45^. Reversely, Head et al.^46^ revealed that mTOR activation is dependent on VDAC1 function. VDACs are considered as regulatory elements of metabolic crosstalk between mitochondria and the cytosol^47^. There are three isoforms: VDAC1, VDAC2 and VDAC3. Whereas VDAC1 and VDAC2 are assumed to share similar functions in regulating the exchange of Ca^2+^, ATP/ ADP and other metabolites, VDAC3 seems to be functionally distinct. Notably, deletion of VDAC2, the isoform TCAIM is interacting with, is the only embryonically lethal one and overexpression of VDAC1 and VDAC3 fail to compensate for VDAC2 KO^47^ highlighting its non-redundant role. Furthermore, it has been described that binding of mTORC1 to VDAC1 and Bcl-xl enhances ΔΨm ^45, 48^.

Interestingly, TCAIM KI CD8^+^ T cells show a reduced activation-induced upregulation of ΔΨm which might be caused by TCAIM-VDAC2 interaction. Our investigations not only revealed delayed MERC formation in TCAIM KI T cells but also an increased length of these late forming subcellular structures. Interestingly, long MERCs have been observed in situations when the mTORC1 nutrient-sensing pathway disengages^49^. Thus, indeed TCAIM interaction with VDAC might interfere with mTORC1 activation at the mitochondria resulting in nutrient-drop-mediated delayed enlargement of MERCs. Furthermore, MERCs and especially the local activity of mTOR and VDAC at MERCs have been previously reported to function as immunometabolic hubs linking TCR signals to enforcement of glycolysis^50^. Subcellular accumulation of mTORC2-AKT-GSK3ȕ at MERCsenabled fast recruitment of hexokinase I to VDAC and thus, induction of metabolic flux into mitochondria. However, this was exclusively shown for rapid memory CD8^+^ T cell responses and limited to glycolytic flux. With our study, we additionally demonstrate an essential role of MERCs and TCAIM for cholesterol biosynthesis and effector cell differentiation during primary T cell activation.

## Methods

### Mice

TCAIM KO mice were purchased from MRC Harwell Institute, C57BL/6N mice from Charles River Laboratories, C57BL/6-Tg(CAG-Flpe)2Arte mice from TaconicArtemis GmbH and Cd4Cre-Car mice were provided by M. Schmitt-Supprian. TCAIM KI Cd4Cre mice were previously described^15^. TCAIM KO mice were crossed first with C57BL/6-Tg(CAG-Flpe)2Arte and subsequent with Cd4Cre-Car mice to obtain TCAIM KO Cd4Cre mice bearing a conditional T cell specific deletion of exon four of the Tcaim gene resulting in a loss of protein function. TCAIM KI Cd4Cre and TCAIM KO Cd4Cre mice strains were on a C57BL/6 background. All mice were bred and/ or housed under specific pathogen-free conditions in the animal facility of the Charité. For experiments, gender-matched male and female littermates or mice from parallel breeding (max. 1 - 7 weeks age divergent) aged between 2 - 6 months were used. Studies were performed in accordance with the guidelines of the Federation of European Laboratory Animal Science Association (FELASA) and approved by the local authority [Landesamt für Gesundheit und Soziales Berlin (LAGeSo)].

### Preparation of single cell suspensions

Spleen and lymph nodes were first forced through 100 µm cell strainers (Falcon/ Corning) with 4 °C cold phosphate buffered saline (PBS, Gibco) and centrifuged at 4 C and 300 g. Then erythrocytes of spleen cell suspensions were lysed in hypotonic PBS diluted in a 1:3 ratio with sterile deionized H2O for 12 seconds. Lysis was stopped by addition of PBS supplemented with 2 % (vol/ vol) fetal calf serum (FCS, Biochrom). Lastly, single cell suspensions of spleen and lymph nodes were filtered through 40 µm cell strainers (Falcon/ Corning) with 4 °C PBS containing 2 % (vol/ vol) FCS. For isolation of colon cells, the entire colon was isolated, rinsed with ice cold Hanks’ Balanced Salt Solution (HBSS) without calcium and magnesium (Gibco) supplemented with 2 % (vol/ vol) FCS and and 10 mM 4-(2-Hydroxyethyl)piperazine-1-ethanesulfonic acid (HEPES, Merck) and cut into 1 cm pieces. Colon pieces were placed in HBSS containing 2 % (vol/ vol) FCS, 10 mM HEPES and 0,154 mg/ ml 1,4-Dithioerythritol (DTE, Sigma) and incubated in a shaker incubator (200 rpm, 37 °C) for 15 minutes. This was repeated twice. Then colon pieces were placed in HBSS containing 2 % (vol/ vol) FCS, 10 mM HEPES and 5 mM Ethylenediaminetetraacetic acid (EDTA, Sigma-Aldrich) and incubated in in a shaker incubator (500 rpm, 37 °C) for 15 minutes. This was repeated twice. Residual EDTA was removed by rinsing colon pieces in VLE RPMI 1640 with glutamine (Biochrom AG). Then colon pieces were digested in RPMI 1640 containing 10 % (vol/ vol) FCS, 0,1 mg/ ml Liberase TL Research Grade (Roche) and 0,1 mg/ ml DNAse I (Roche) for 20 minutes in a shaker incubator (200 rpm, 37 °C). Following incubation, remaining tissue pieces were first forced through 100 µm cell strainers with HBSS containing 10 mM HEPES and then filtered through 40 µm cell strainers with VLE RPMI 1640 with glutamine containing 10 % (vol/ vol) FCS, 100 U/ mL penicillin and 100 mg/ mL streptomycin (all Biochrom AG).

### Cell culture and cell lines

Naive CD8^+^ T cells were isolated from single cell suspensions of spleen and lymph nodes using a Naive CD8^+^ T Cell Isolation Kit, mouse (Miltenyi) or EasySep Mouse Naïve CD8+ T Cell Isolation Kit (StemCell) and cultured in VLE RPMI 1640 with glutamine supplemented with 10 % (vol/ vol) FCS, 100 U/ mL penicillin, 100 mg/ mL streptomycin (all Biochrom AG) and 50 µM ȕ-mercaptoethanol (Sigma-Aldrich) at 37 °C and 5 % CO2. If not stated otherwise, 1 x 10^5^ cells were polyclonally stimulated with 2 µg/ ml plate-bound αCD3 (145-2C11, eBioscience) and 1 µg/ ml soluble αCD28 (37.51, eBioscience) for the indicated time. To quantify CD8^+^ T cell proliferation, cells were stained with Cell Proliferation Dye eFluor™ 450 (eBioscience) according to manufacturer’s protocol prior culture. If indicated, cyclosporin A (Merck), DMSO (Sigma-Aldrich) or cholesterol–methyl-ȕ-cyclodextrin (Sigma-Aldrich) was added to the culture in increasing concentrations (0.01 – 0.5 µM, 10 µM, or 2 – 40 µM, respectively). For some experiments, naïve CD8^+^

T cells had been co-cultured with autologous B cells isolated from spleen cell suspensions using an EasySep Mouse B Cell Isolation Kit (StemCell) in a 1:1 ratio. HEK293T cell line (ACC 305) was purchased from the DSMZ-German Collection of Microorganisms and Cell Cultures GmbH and maintained in DMEM (very low endotoxin, Biochrom AG and PAN Biotech) supplemented with 10 % (vol/ vol) FCS, 100 U/ mL penicillin, 100 mg/ mL streptomycin at 37 °C and 5 % CO2.

### Non-targeted metabolite analysis

Sample preparation and GC/APCI/MS (gas chromatographic atmospheric pressure chemical ionization mass spectrometry) analysis was performed as described previously^51^. Briefly, metabolites of pellets of 7 x 10^5^ 60 h polyclonally activated CD8^+^ T cells and 10 µl of supernatants from 0, 12, 20, 36, 44 and 60 h CD8^+^ T cell cultures were extracted by incubation with 90 µl ice cold methanol for 30 min under shaking at 4 °C. Subsequently, 50 µl H2O was added, samples were vortexed and centrifuged at 21,000 g for 5 min at room temperature. Supernatants were vacuum dried and stored at -80 °C until measurement. For measurement, the dried mixtures were derivatized on-line using a PAL RTC autosampler (CTC Analytics). After addition of 10 μl methoxyamine (20 mg/ ml in pyridine; Sigma), vials were agitated for 90 min at 34 °C and 750 rpm. 90 μl N-methyl-N-(trimethylsilyl)-trifluoroacetamide (MSTFA; Macherey-Nagel) containing 0.2 μg/ ml each of C4-C24 fatty acid methyl esters (Sigma) as retention index markers were added, followed by agitating vials for 30 min. Before injection, samples were allowed to rest for 2 h to complete derivatization reactions. GC-APCI-MS analysis was carried out on an Agilent 7890 B gas chromatograph (Agilent) coupled to an Impact II quadrupole time-of-flight (Q-TOF) mass spectrometer (Bruker) via a GC-APCI II source (Bruker). 1 μl sample was injected into a split/ splitless inlet, operated at 230 °C in split mode (1:10). Chromatographic separation was carried out on a 30 m x 0.25 mm x 0.25 μm HP5-MS UltraInert capillary column (Agilent) connected to a 0.5 m x 0.25 mm RxiGuard (Restek) column as transfer capillary. Helium was used as a carrier gas at 1 ml/ min. The temperature gradient was as follows: 80 °C held for 2 min, 15 °C/ min to 320 °C, 320 °C held for 7 min. Transfer capillary and capillary head of the GC-APCI II source were heated to 290 °C. The GC-APCI source was operated in positive ion mode, using a capillary voltage of -2500 V and an end plate offset of -250 V. A corona current of 3000 nA and a nebulizer pressure of 2.0 bar was used. The dry gas was nitrogen at a flow rate of 2.0 l/ min and a temperature of 230 °C. Full-scan line spectra were recorded in the scan range of m/ z 80 - 1000 at an acquisition rate of 10 s^-1^. Q-TOF ion transfer parameters were as follows: funnel 1 RF 300.0 Vpp, funnel 2 RF 300.0 Vpp, isCID energy 0.0 eV, hexapole RF 100.0 Vpp, quadrupole ion energy 4.0 eV, low mass 80.0 m/ z, collision energy 8.0 eV, collision RF 1000.0 Vpp, transfer time 75 μs, pre-pulse storage 8.0 μs. External mass calibration was performed before each sequence by direct infusion of 180 μl/ h ES low concentration tune mix (Agilent) according to manufacturer’s instruction. Subsequently, metabolites were identified in comparison to the Golm Metabolome Database^52^ or putatively annotated as described previously^53^. Following identification, metabolite peak intensities were log10 transformed and normalized for experimental replication and run order effects using an ANOVA based method^54^.

Log-transformation improves normality of the data distribution. Run order effects are systematic effects on measured metabolite intensities due to the measurement time point.

Experimental replications describe the samples being generated in three independent laboratory trials. ANOVA based normalization removes effects related to these factors prior to further statistical tests.

### Flow cytometry

Dead cells were stained using either Zombie UV™ Fixable Viability Kit (Biolegend, 1:1000), Fixable Viability Dye eFluor™ 506 (eBioscience, 1:100) or LIVE/DEAD™ Fixable Far Red Dead Cell Stain Kit (Invitrogen, 1:5000) according to manufacturer’s protocol. Concomitantly, Fc receptors were blocked with Purified Rat Anti-Mouse CD16/CD32 Mouse BD Fc Block™ (2.4G2, BD Pharmingen, 1:1000). Cell surface antigens were stained in PBS containing 2 % (vol/ vol) FCS and 1 mg/ ml sodium azide (Serva) at 4 °C in the dark. Cell fixation and staining of intracellular antigens was done with Foxp3 Staining Buffer Set (Miltenyi) according to manufacturer’s protocol. For cytokine detection, cells were treated prior staining with 10 ng/ mL phorbol 12-myristate 13-acetate (Sigma-Aldrich) and 1 µg/ mL ionomycin (Biotrend) for 4 h at 37 °C. For the last 2 h 2 µg/ mL Brefeldin A (Sigma-Aldrich) was added. Monoclonal antibodies against the following antigens were used: CD3 (17A2, Biolegend, CD8a (53-6.7, Biolegend or BD Pharmingen), CD25 (PC61, Biolegend), CD44 (IM7, Biolegend), CD45R (RA3-6B2, Biolegend or Miltenyi),

CD62L (MEL-14, Biolegend), CD69 (H1.2F3, Biolegend), GZMB (NGZB, eBioscience), Hif1α (Mgc3, eBioscience), IFN-Ȗ (XMG1.2, Biolegend), Ki67 (SolA15, eBioscience)), TCRȕ (H57-597, Biolegend). For measurement of ΔΨm or ROS production, cells were stained with 0.1 µM MitoStatus TMRE (BD Pharmingen) for 20 min or 5 µM MitoSOX™ Red mitochondrial superoxide indicator (Invitrogen) for 10 min at 37 °C according to manufacturer’s protocol, respectively, and subsequently, stained for dead cells and surface antigens simultaneously in PBS or HBSS with Ca^2+^/ Mg^2+^ (Gibco) for 10 min at 4 °C. Data were acquired using an LSRFortessa with BD FACSDiva software v8.0.2 (BD Biosciences) or a CytoFLEX LX with CytExpert software v2.3.1.22 (Beckman Coulter) and analyzed with FlowJo v10 software (FlowJo, LLC).

### Immunohistology

Colon samples were fixed in 36.5 % formaldehyde and embedded in paraffin. Sections of 1 – 2 µm thickness were cut, deparaffinated and subjected to a heat-induced epitope retrieval step. Endogenous peroxidase was blocked by hydrogen peroxide prior to incubation with αCD3 (polyclonal rabbit, Agilent #IR50361-2) followed by the EnVision+ System-HRP Labelled Polymer Anti-Rabbit (Agilent). For visualization, OPAL-570 diluted in amplifier diluent (Akoya Biosciences) was used. Proteins were then inactivated and sections incubated with αCD4 (4SM95, eBioscience) followed by incubation with rabbit anti-rat secondary antibody (Dianova), then with the EnVision+ System-HRP Labelled Polymer Anti-Rabbit and finally with OPAL-670 diluted in amplifier diluent (Akoya Biosciences). The procedure was repeated for αCD8 (4SM15, eBioscience) staining using OPAL-520 (Akoya Biosciences) for visualization. Nuclei were stained using 4′,6-diamidine-2′-phenylindole dihydrochloride (DAPI, Sigma). Slides were coverslipped in Fluoromount G (Southern Biotech). Multispectral images were acquired using a Vectra® 3 imaging system employing the Vectra3 and Phenochart software (all Akoya Biosciences). Images are displayed in pseudocolours. Cells were quantified in a blinded manner using the inForm Software (Akoya Biosciences).

### Quantitative real time PCR

RNA of polyclonally activated CD8^+^ T cells from C57BL6/N mice was purified using NucleoSpin RNA kit (Macherey-Nagel) according to manufacturer’s protocol. RNA was then incubated at 65 °C for 5 min. Subsequently, cDNA was synthesized using QuantiTect Reverse Transcription kit (Qiagen). Gene expression was quantified using TaqMan gene expression assays (Thermo Fisher) for Tcaim and Ubc or custom designed primer and probes for Il-2 and Ifn-*γ* (Eurofins Genomics) and TaqMan Universal PCR Master Mix (Thermo Fisher) on a 7500 Real Time PCR System (Applied Biosystems). Sequences for murine Il-2: forward primer: TCGCCAGTCAAGAGCTTCAGACAAGCA, reverse primer: CATGCCGCAGAGGTCCAA, probe: CAATTCTGTGGCCTGCTTGGGCAA. Sequences for murine Ifn-*γ*: forward primer: AGCAACAGCAAGGCGAAAAA, reverse primer: AGCTCATTGAATGCTTGGCG, probe: ATTGCCAAGTTTGAGGTCAACAACCCACA.

Probes were coupled to FAM and TAMRA. The thermal cycling program was set as followed: 2 min at 50 °C, 10 min at 95 °C and 40 cycles of 15 s at 95 °C and 1 min at 60 °C. The reactions were performed as duplicates. Data were analyzed with 7500 Systems SDS Software v2.3 and relative gene expression was calculated as 2^-ΔCT^ using Ubc as reference gene.

### RNA-seq

RNA of naïve and polyclonally activated CD8^+^ T cells was purified using NucleoSpin RNA kit (Macherey-Nagel) according to manufacturer’s protocol and submitted to the Scientific Genomics Platform (Max Delbrück Center for Molecular Medicine) for sequencing. Briefly, mRNA-Seq libraries were prepared from 100 ng of input total RNA samples using TruSeq stranded mRNA kit (Illumina). Barcoded and pooled libraries were sequenced on 5.5 lanes of Illumina HiSeq 4000 platform (HCS v3.3.76, RTA v2.7.6) in a paired-end 2 x 75 nt sequencing mode. Fastq files generation and demultiplexing were performed using bcl2fastq v2.20.0.

### RNA-seq raw data processing

The quality of fastq files was controlled by FastQC v0.11.8 (Bioinformatics Group at the Babraham Institute). Residual adapter sequences, ambiguous bases, low quality reads and reads with a length below 25 were removed by AdapterRemoval v2.2.3^55^. The remaining reads were aligned to the GRCm38 Ensembl version of the mm10 mouse genome using Tophat v2.1.1^56^ and Bowtie2 v2.2.5^57^. Counts per gene were summarized by the featureCount algorithm of the Rsubread package v2.4.2^58^ in R v4.0.2 (R Core Team (2020). R: A language and environment for statistical computing. R Foundation for Statistical Computing, Vienna, Austria. URL https://www.R-project.org/.).

### RNA-seq data analysis

Raw count data of protein-coding genes were normalized and either log2-transformed after adding a pseudocount to each value or variance-stabilized transformed using the DESeq2 package v1.30.0^59^ in R. 1000 genes with the highest variances across all samples were selected and z-scores of the normalized expression values of these genes were subjected to hierarchical clustering. The heatmap representation of the clustering result was done with the in-built heatmap function in R. Z-scores of expression values of further genes preselected by functional aspects were shown in a heatmap with predefined order created with the pheatmap package v1.0.12 (Raivo Kolde (2019). pheatmap: Pretty Heatmaps. R package version 1.0.12. https://CRAN.R-project.org/package=pheatmap).

Differential expressed genes between two groups were determined by fitting models of negative binomial distributions to the count data. Raw p-values were adjusted for multiple testing using false discovery rate. Genes with significant differential expression were selected by an adjusted p-value below 0.05 and a minimal absolute log2-fold change of one. Enrichment of functional aspects within the sets of differential expressed genes were analyzed after mapping the genes to functions by DAVID using the R-packages RDAVIDWebService v1.26.0^60^ and clusterProfiler v3.16.1^61^. Selected results were plotted using GOCircle - function of the GOplot package v1.0.2^62^.

### Mathematical metabolite flux modelling

We modeled glycolysis, serine biosynthesis, TCA cycle, and glutaminolysis pathways as they represent the main carbon flux routes for biomass precursors and energy production, therefore, reflecting the major changes in metabolic activity. The overall network was constructed to have at maximum two unmeasured intermediate metabolites except in reactions from glycolysis or TCA cycle. All carbon molecules taken up were either converted into biomass building blocks or secreted. To estimate the biomass of murine CD8^+^ T cells, we used an adapted version of Thiele’s human lymph node germinal center biomass reaction as the closest available genome scale metabolic reconstruction^63^. Growth based dilution was negligible in this model as the fluxes were orders of magnitude higher than the dilution fluxes. The relative measurements from the non-targeted metabolite analysis of TCAIM KI and wt CD8^+^ T cells and their culture supernatants were converted to absolute values using calibration data from Lisec et al.^64^ combined with log-log-regression. The kinetics of the model were either reversible or irreversible mass action kinetics. The kinetic rates of TCAIM KI CD8^+^ T cells were considered as deviation from the kinetic rates of wt CD8^+^ T cells and, therefore, expanded with regulation parameters. All co-factors were held constant at a value of 1 mM as they were not measured in the experiments. We assumed steady state for the entire model. The objective of the optimization was the minimization of the sum of the absolute values of the logarithm of regulation parameters, representing a minimal metabolic change between wt and TCAIM KI CD8^+^ T cells. We further minimized the sum of glutamine uptake for wt and TCAIM KI CD8^+^

T cells together with the sum of regulation parameters to avoid non-physiological futile cycling in the TCA cycle. To optimize the objective function, we used the R package “nloptr” (Steven G. Johnson, The NLopt nonlinear-optimization package, http://github.com/stevengj/nlopt) in a multi start approach to avoid local optima. Further details of the modelling methods are described in (Identification of differential regulation in central carbon metabolism between related cell lines, Roman Rainer, 2020, Humboldt-Universität zu Berlin, http://dx.doi.org/10.18452/22117).

### Plasmids and plasmid constructions

The p TCAIM-EGFP-N1 construct was generated as previously described^16^ (TCAIM was formerly known as TOAG1). Mito-PAGFP encoding for an eGFP with mitochondrial target sequence was a gift from Richard Youle (Addgene plasmid # 23348; http://n2t.net/addgene:23348; RRID:Addgene_23348)^65^.

For split eGFP fusion protein generation, Tcaim/ Tomm40/ Immt sequences containing a 3’- end SapI restriction site inserted into pMA-T vectors as well as c- and n-terminal parts of split eGFP sequences inserted into pMA-RQ and pMK-RQ, respectively, were purchased from Invitrogen. C- and n-terminal split eGFP fragments were ligated into TCAIM, TOM40 or MIC60 coding pMA-T vectors by destroying Tcaim/ Tomm40/ Immt stop codons to create the respective split GFP fusion proteins. Fusion protein coding sequences were excised by BamHI and NotI restriction and inserted into BamHI and NotI restricted pRNA2 vector (gift from Prof. Nina Babel) for expression analysis in eukaryotic systems.

### Co-immunoprecipitation and mass spectrometry analysis

1.5 x 10^6^ HEK293T cells were seeded in DMEM supplemented with 10 % (vol/ vol) FCS, 100 U/ mL penicillin, 100 mg/ mL streptomycin on a 10 cm tissue culture treated petri dish (Falcon) and 24 h later transfected with either p TCAIM-EGFP-N1 (25 µg) or mito-PAGFP (10 µg) control plasmid DNA using the calcium phosphate technique. In brief, calcium-phosphate-DNA-particle were generated by mixing the plasmid DNA’s with a 250 mM CaCl (Sigma-Aldrich) solution which was then added dropwise to an equal volume of 2 x hepes buffered saline and subsequently incubated for 20 min at room temperature. The precipitated particles were then added dropwise to the cells. After 24 h of incubation at 37 °C and 5 % CO2 the medium was replaced for fresh supplemented DMEM culture medium. The co-immunoprecipitation was then performed utilizing the µMACS GFP Isolation Kit (Miltenyi) with protein extracts from whole cell lysates of 2 x 10^7^ transfected HEK293T cells. Deviating to manufacturer’s protocol cells were incubated in lysis buffer supplemented with protease inhibitor (Sigma-Aldrich) in a 1:10 ratio for 1 h on ice. Eluates were separated by SDS-PAGE, stained with PageBlue (Thermo Scientific) and analyzed by liquid chromatography mass spectrometry (LC-MS/MS). Briefly, protein bands in molecular weight range of 40 – 250 kDa were excised and subjected to in-gel tryptic digestion as described previously ^66^. LC-MS/MS analyses were performed on an Ultimate 3000 RSLCnano system online coupled to an Orbitrap Q Excative Plus mass spectrometer (both Thermo Fisher Scientific). The system comprised a 75 µm i.d. × 250 mm nano LC column (Acclaim PepMap C18, 2 µm; 100 Å; Thermo Fisher Scientific). Mobile phase (A) was 0.1 % formic acid in water and (B) 0.1 % formic acid in 80:20 (v/v) acetonitrile/water. The gradient was 8 – 38 % B in 80 min. The Q ExactivePlus instrument was operated in data dependent mode to automatically switch between full scan MS and MS/MS acquisition. Survey full scan MS spectra (m/ z 350 – 1650) were acquired in the Orbitrap with 70,000 resolution (m/ z 200) after 60 ms accumulation of ions to a 1 x 10^6^ target value. Dynamic exclusion was set to 60 s. The ten most intense multiply charged ions (z ≥ 2) were sequentially isolated and fragmented by higher-energy collisional dissociation (HCD) with a maximal injection time of 120 ms, AGC 5 x 10^4^ and resolution 17,500.

For protein identification and relative label-free quantification, MaxQuant software v1.6.0.1^67^ with default Andromeda LFQ parameter was used. Spectra were matched to a mus musculus (17,040 reviewed entries, downloaded from uniprot.org), a contaminant and decoy database. Acquired MS/MS were searched with parameter as followed: precursor mass tolerance of 10 ppm, fragment tolerance of 0.02 Da, trypsin specificity with a maximum of two missed cleavages, cysteine carbamidomethylation set as fixed and methionine oxidation as variable modification. Identifications were filtered at 1 % False Discovery Rate (FDR) at peptide level. Bioinformatic analysis of the data was performed with Perseus software v1.6.0.2.

### BiFC Assay

4 x 10^4^ HEK293T cells were seeded in DMEM supplemented with 10 % (vol/ vol) FCS, 100 U/ mL penicillin, 100 mg/ mL streptomycin on a tissue culture treated 24 well plate (Falcon) and 24 h later co-transfected with either p TCAIM-EGFP^C^-RNA2 and pTOM40-EGFP^N^-RNA2, p TOM40-EGFP^N^-RNA2 and pMIC60-EGFP^C^-RNA2 or pTCAIM-EGFP^C^-RNA2 alone using Lipofectamine 2000 Transfection Reagent (Invitrogen) according to manufacturer’s protocol. A total volume of 500 ng DNA (250 ng per construct) and a final volume of 4 µl Lipofectamine was used for each transfection. After 72 h of incubation at 37 °C and 5 % CO2 cells were trypsinized using Trypsin-EDTA (PAN Biotech) diluted 1:2 in PBS, washed and eGFP expression was examined by flow cytometry using an LSRFortessa with BD FACSDiva software v8.0.2 (BD Biosciences). Data were analyzed with FlowJo v10 software (FlowJo, LLC).

### Transmission electron microscopy

Naïve and polyclonally activated CD8^+^ T cells were fixed with 0.1 M sodium cacodylate buffer (Serva) supplemented with 2.5 % glutaraldehyde (Serva) for 30 min at room temperature, stored by 4 °C and submitted to the Core Facility for Electron Microscopy (Charité – Universitätsmedizin Berlin). Samples were postfixed with 0.1 M sodium cacodylate buffer supplemented with 1 % osmium tetroxide (Science Services) and potassium ferrocyanide (Merck) and embedded in 1 % agarose (Sigma-Aldrich). Embedded samples were dehydrated in a graded ethanol (Merck) series and embedded in epon (Serva). Ultrathin sections of 70 nm were prepared on a Leica Ultracut S (Leica Biosystems). Sections were stained with 4 % uranyl acetate (Serva) and Reynold’s lead citrate (Merck). Micrographs were taken with a Zeiss EM 906 transmission electron microscope at 80 kV acceleration (Carl Zeiss) and a slow scan 2k CCD camera (TRS). Mitochondria and cristae were identified manually and the measures were outlined with an optical pen using ImageJ v1.53a software. The roundness factor was calculated as 4*area/pi*sqr(major axis) and a maximal distance of 75 nm between mitochondria and ER was defined as MERC.

### Transcription factor binding site prediction

A genome sequence of ∼900 bp in close proximity to the transcriptional start site (TSS) and upstream of the translational start (ATG) of the murine Tcaim gene was selected via Ensembl genome browser analyzed using the TFSEARCH online tool (http://diyhpl.us/~bryan/irc/protocol-online/protocol-cache/TFSEARCH.html) with a statistical score threshold above 85 %^68^.

### Statistical analysis

Quantitative data were represented as scatter dot plots, means, means ± SEM, boxplots with whiskers from min to max or stacked bar charts as indicated in the figure legends. Except for electron microscopy, graph data points of scatter dot plots and boxplots with whiskers from min to max represent biological replicates. Statistical significance was determined as indicated in the figure legends. Datasets were first tested for Gaussian distribution using the D’Agostino and Pearson normality test for sample sizes above five. For smaller sample sizes, the Shapiro-Wilk normality test was applied. If both compared datasets were Gaussian distributed a parametric test was applied, otherwise a non-parametric test was used. P-Values below 0.05 were considered as statistically significant. Statistical analysis was done using GraphPad Prism v7 software (GraphPad Software).

## Data availability

The RNA-seq data are deposited at the NCBI GEO database under the accession code: GSE167520.

## Acknowledgements

We thank the Core Facility for Electron Microscopy of the Charité, (especially Petra Schrade) for support in acquisition of the data. We also thank Dr. T. Borodina and M. Sohn from the Scientific Genomics Platform (Max Delbrück Center for Molecular Medicine) for library preparation and sequencing

## Author contributions

C.I., J.S. and B.S. designed the experiments. C.I. performed the experiments with the assistance from C.A., K.V., K.M., K.S., and D.U., analyzed the data and performed the statistical analysis with the exception of: J.L. performed the GC/APCI/MS and analyzed the metabolomics data; A.A.K. performed the immunohistology and analyzed the data; K.J. analyzed the RNA-seq data; R.J.R. constructed the metabolic flux models with the assistance and advice of K.T. and E.K; K.J., K.T.T. and C.G. performed the LC-MS/MS and identification of TCAIM interaction partner. A.P. and S.S. contributed critical reagents and provided experimental help or helped with data analysis. C.M. provided advice to data interpretation. C.I. and B.S. wrote the manuscript.

---

**Extended Data Fig. 1.**
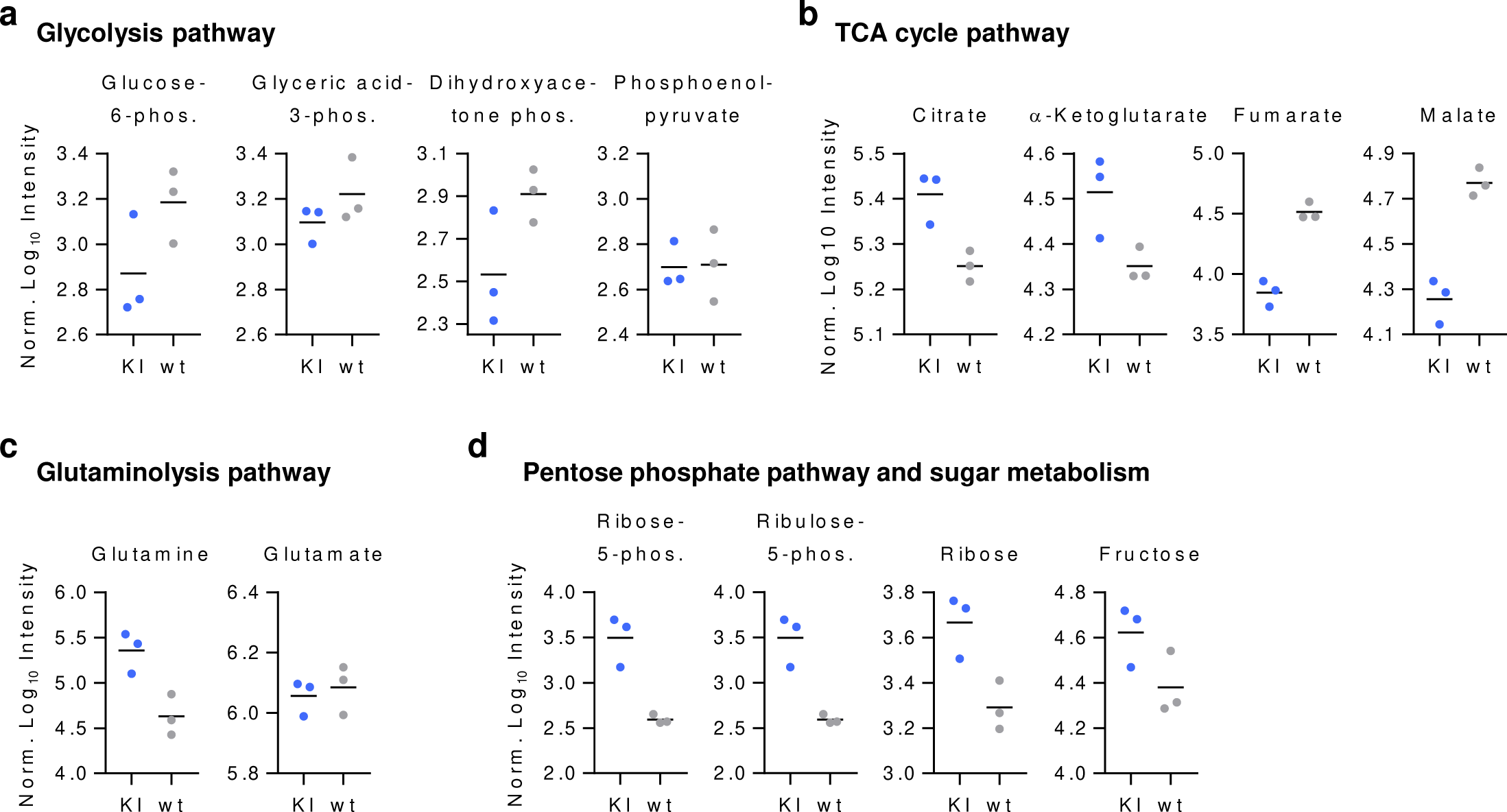
TCAIM KI causes an altered metabolic program in activated CD8^+^ T cells. Non-targeted metabolite analysis identified by GC/APCI mass spectrometry of 60 h polyclonally activated TCAIM KI or wt CD8^+^ T cells (n = 3). Cells had been stimulated with 5 µg/ ml plate-bound αCD3 and 2 µg/ ml soluble αCD28 at a density of 6 x 10^5^ cells/ 200 µl culture medium. Selected metabolites of the glycolysis (**a**), TCA cycle (**b**), glutaminolysis (**c**), as well as pentose phosphate and sugar metabolism (**d**) pathway are shown. Quantitative data represents means.

**Extended Data Fig. 2.**
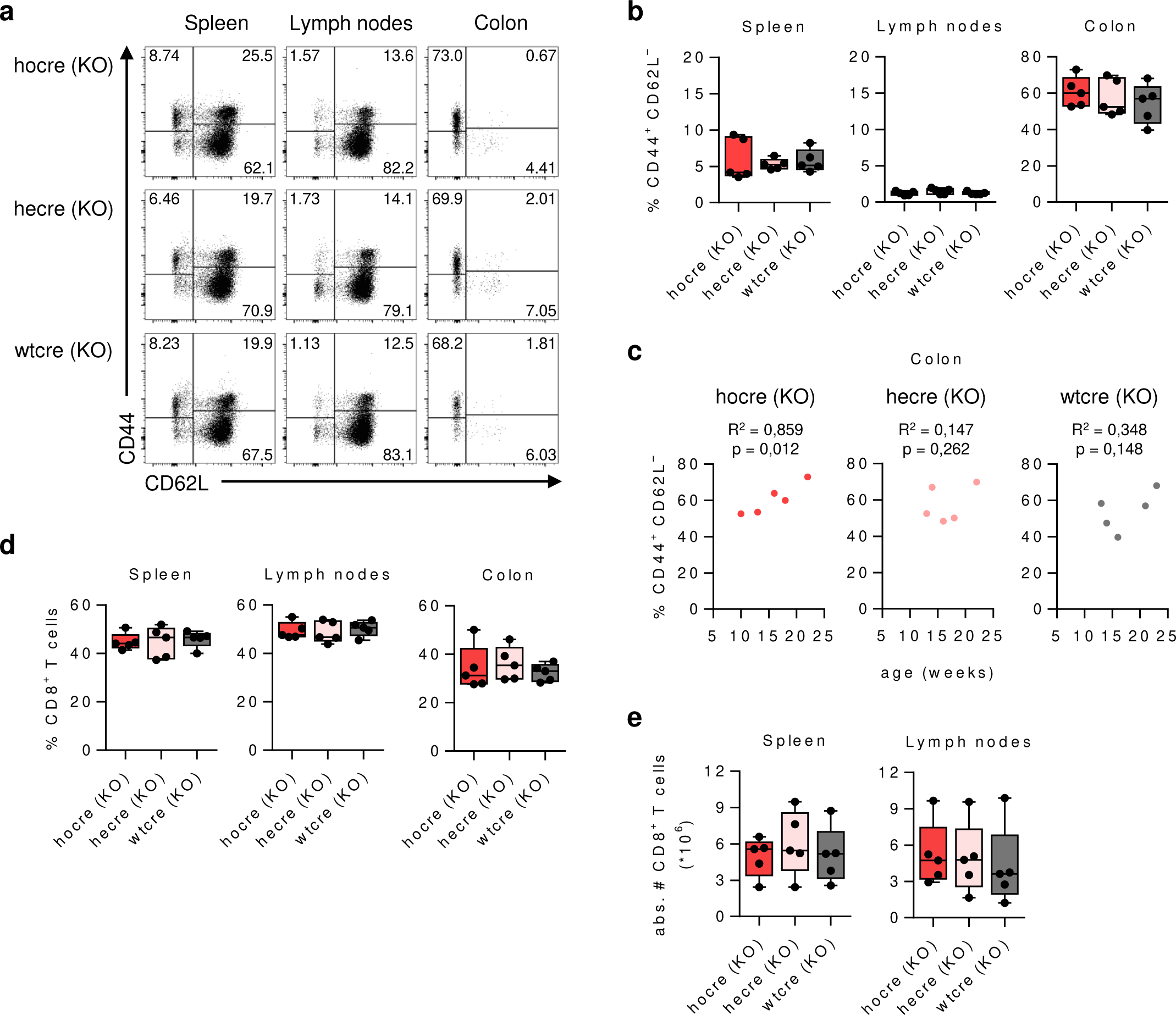
TCAIM KO favors spontaneous effector differentiation of peripheral CD8^+^ T cells. Frequencies (**a**) and absolute numbers (**b**) of CD8^+^ T cells residing in the spleen, lymph node or colon of mice with homozygous (hocre), heterozygous (hecre) TCAIM KO or wild type (wtcre) gene expression measured ex vivo by flow cytometry. Representative dot plots (**c**) and quantification of CD62L– / CD44+ effector/ effector memory CD8⁺ T cell frequencies (d) from TCAIM KOCd4Cre mice with the indicated genotype and tissue. (**e**) Correlation analysis between the mice’s age and the frequency of colon residing CD62L– / CD44+ effector/ effector memory CD8⁺ T cells from TCAIM KOCd4Cre mice with homozygous (red), heterozygous (pink) TCAIM KO or wild type (dark grey) gene expression. Squared Pearson’s correlation coeficient (R^2^) and p-values are shown for each one-tailed comparison. Quantitative data represents boxplots with whiskers from min to max (n = 5).

**Extended Data Fig. 3.**
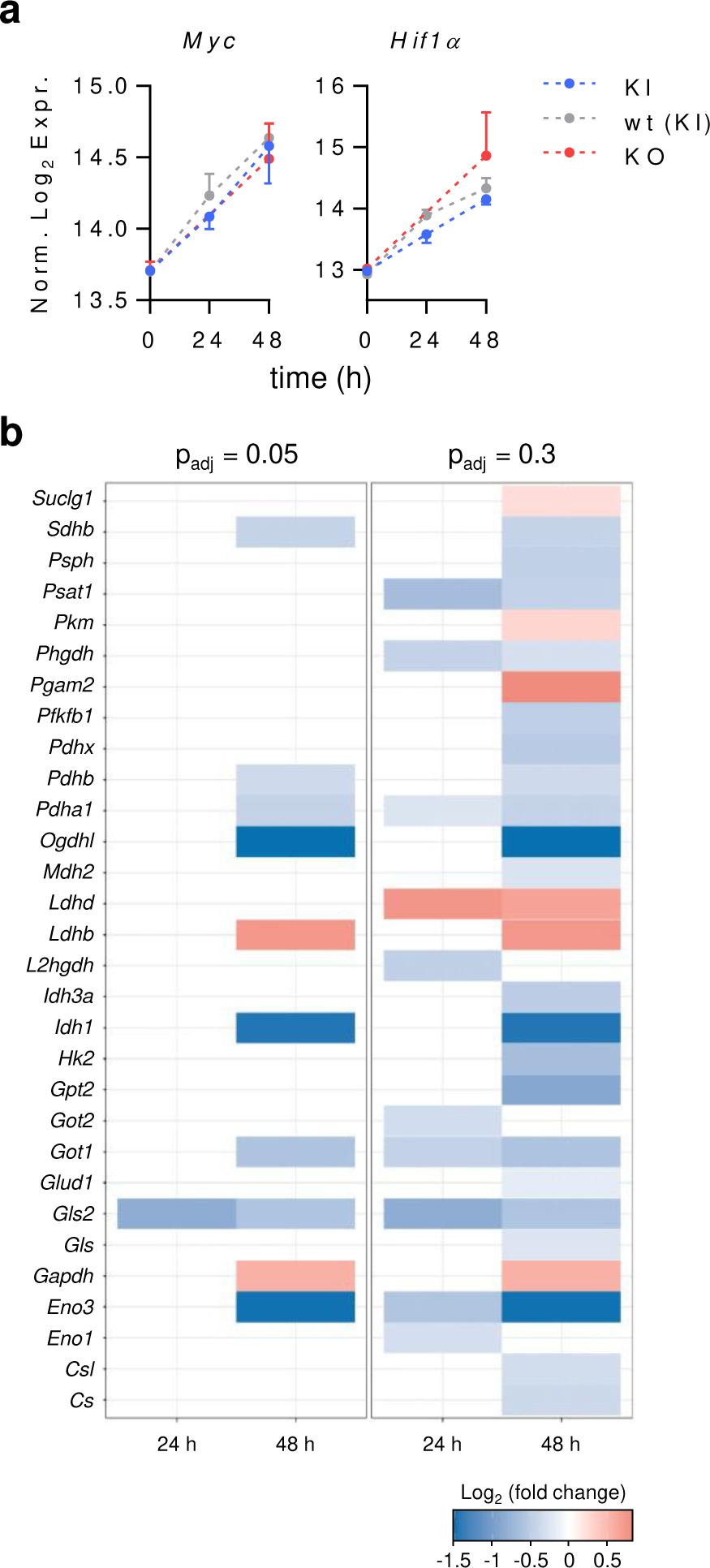
TCAIM-dependent expression of metabolism controlling genes in activated CD8^+^ T cells. (**a**) Myc and *Hif1α* mRNA expression of naïve and polyclonally activated CD8^+^ T cells with TCAIM KI, KO or wt gene expression (KI and wt: n = 3; KO: n = 2). RNA-seq raw count data of protein-coding genes were normalized and log_2_-transformed after adding a pseudocount to each value. Cells had been stimulated with 5 µg/ ml plate-bound αCD3 and 2 µg/ ml soluble αCD28 at a density of 6 x 10^5^ cells/ 200 µl culture medium. Quantitative data represents means ± SEM. (**b**) Overview of the differentially expressed enzymes controlling the glycolysis, TCA cycle and glutaminolysis pathway between 24 to 48 h polyclonally activated TCAIM KI and wt CD8^+^ T cells (log_2_ fold change). P-values have been adjusted to either 0.05 or 0.3 as indicated (n = 3). White areas indicate p-values exceeding the adjusted ones. Cells had been stimulated with 5 µg/ ml plate-bound αCD3 and 2 µg/ ml soluble αCD28 at a density of 6 x 10^5^ cells/ 200 µl culture medium (a-b).

**Extended Data Fig. 4.**
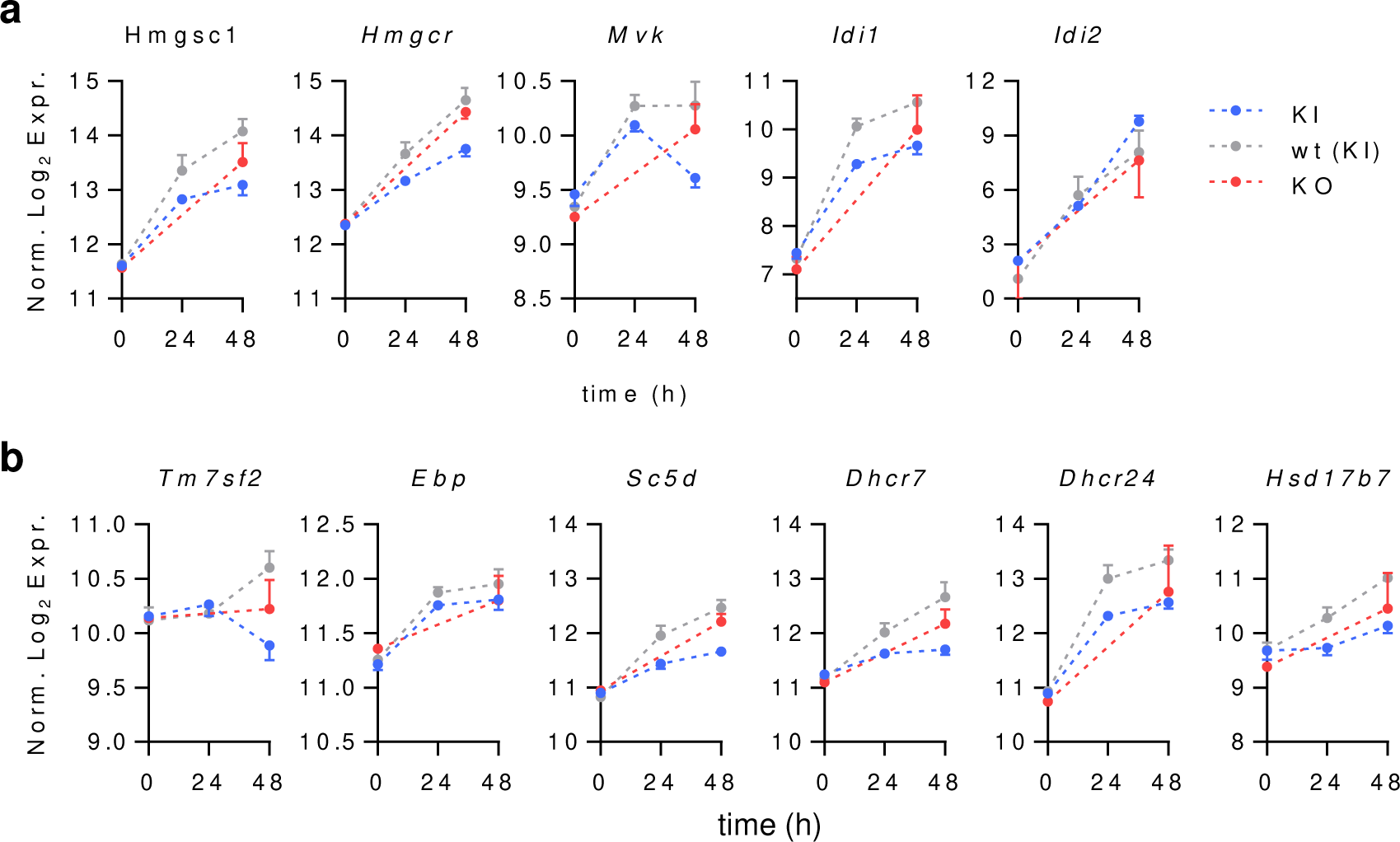
Reduced gene expression of mevalonate pathway and cholesterol biosynthesis controlling enzymes in TCAIM KI CD8^+^ T cells. mRNA expression of additional genes controlling the mevalonate pathway (**a**) and the cholesterol biosynthesis (**b**) of naïve and polyclonally activated CD8^+^ T cells with TCAIM KI, KO or wt gene expression (KI and wt: n = 3; KO: n = 2). RNA-seq raw count data of protein-coding genes were normalized and log_2_-transformed after adding a pseudocount to each value. Cells had been stimulated with 5 µg/ ml plate-bound αCD3 and 2 µg/ ml soluble αCD28 at a density of 6 x 10^5^ cells/ 200 µl culture medium. Data represents means ± SEM.

**Extended Data Fig. 5.**
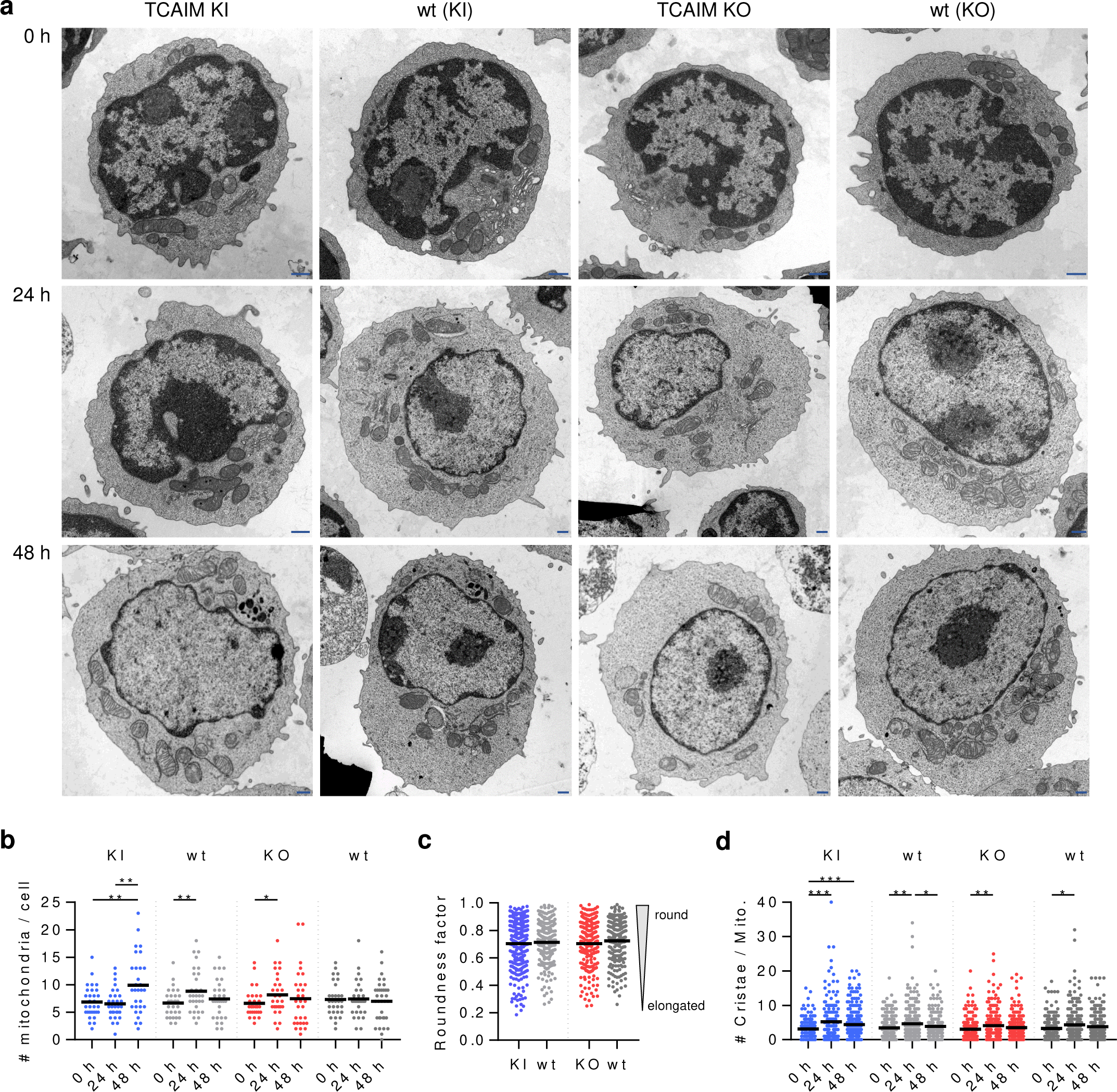
CD8^+^ T cells display an activation-dependent increase in number of mitochondria and cristae. (**a**) Representative transmission electron microscopy images of naïve (upper row) and 24 to 48 h (middle and bottom row, respectively) polyclonally activated CD8^+^ T cells with TCAIM KI, KO or respective to mouse strain wt gene expression. Scale bars: 500 nm. (**b-d**) Quantitative plots of transmission electron microscopy image analysis of naïve and polyclonally activated TCAIM KI, KO and respective wt CD8^+^ T cells. Absolute number of mitochondria per cell (**b**), mitochondrial roundness factor of 48 h activated CD8^+^ T cells only (**c**) and absolute number of cristae per mitochondrion (**d**). For each, mitochondria of 30 cells have been analyzed. Data are represented as scatter dot plots and were analyzed in accordance to Gaussian distribution of the compared sample sets by Kruskal-Wallis test with Dunn’s multiple comparisons test and subsequent post hoc one-tailed, unpaired Student’s t-test (b: KI 0 h vs. 48 h and 24 h vs. 48 h, wt 0 h vs. 24 h) or one-tailed, unpaired Mann-Whitney test (b: KO 0 h vs. 24 h; c: all). * p < 0.05, ** p < 0.01, *** p < 0.001.

**Extended Data Table 1.**
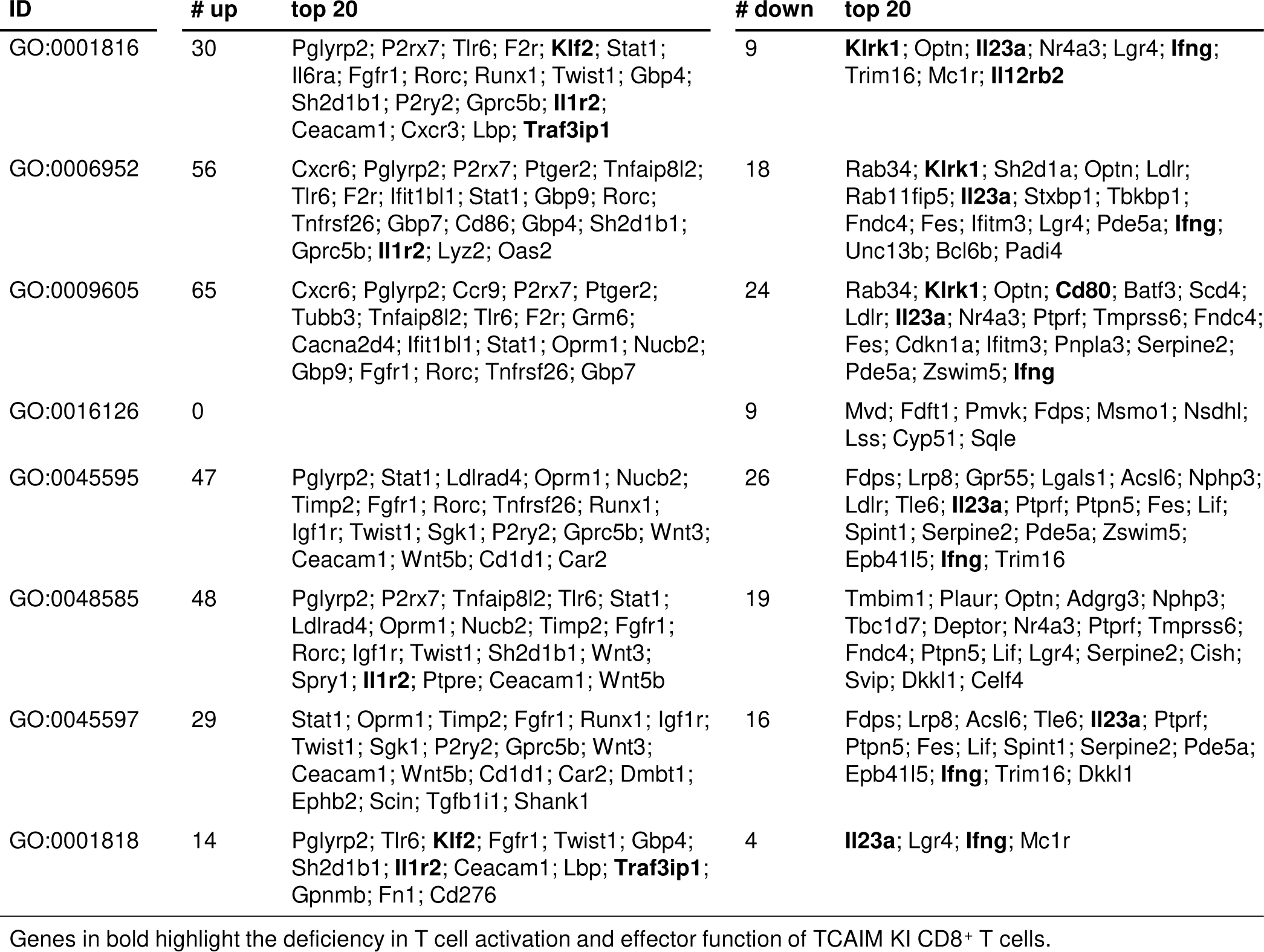
Differentially expressed genes being up- or downregulated in 48 h activated TCAIM KI compared to wt CD8^+^ T cells for chosen annotated GO terms.

## References

1. Shyer, J.A., Flavell, R.A. & Bailis, W. Metabolic signaling in T cells. Cell Res. 30, 649–659 (2020).

2. Chang, C.H. et al. Posttranscriptional control of T cell effector function by aerobic glycolysis. Cell 153, 1239–1251 (2013).

3. Wang, R.N. et al. The Transcription Factor Myc Controls Metabolic Reprogramming upon T Lymphocyte Activation. Immunity 35, 871–882 (2011).

4. Finlay, D.K. et al. PDK1 regulation of mTOR and hypoxia-inducible factor 1 integrate metabolism and migration of CD8(+) T cells. J. Exp. Med. 209, 2441–2453 (2012).

5. Doedens, A.L. et al. Hypoxia-inducible factors enhance the effector responses of CD8(+) T cells to persistent antigen. Nat. Immunol. 14, 1173–U1199 (2013).

6. Angajala, A. et al. Diverse Roles of Mitochondria in Immune Responses: Novel Insights Into Immuno-Metabolism. Front. Immunol. 9 (2018).

7. Cao, Y., Rathmell, J.C. & Macintyre, A.N. Metabolic Reprogramming towards Aerobic Glycolysis Correlates with Greater Proliferative Ability and Resistance to Metabolic Inhibition in CD8 versus CD4 T Cells. PLoS One 9 (2014).

8. Sukumar, M. et al. Mitochondrial Membrane Potential Identifies Cells with Enhanced Stemness for Cellular Therapy. Cell Metab. 23, 63–76 (2016).

9. Murphy, M.P. & Siegel, R.M. Mitochondrial ROS Fire Up T Cell Activation. Immunity 38, 201–202 (2013).

10. Okoye, I. et al. The protein LEM promotes CD8(+) T cell immunity through effects on mitochondrial respiration. Science 348, 995–1001 (2015).

11. Sena, L.A. et al. Mitochondria Are Required for Antigen-Specific T Cell Activation through Reactive Oxygen Species Signaling. Immunity 38, 225–236 (2013).

12. Yerinde, C., Siegmund, B., Glauben, R. & Weidinger, C. Metabolic Control of Epigenetics and Its Role in CD8(+) T Cell Differentiation and Function. Front. Immunol. 10, 2718 (2019).

13. Buck, M.D. et al. Mitochondrial Dynamics Controls T Cell Fate through Metabolic Programming. Cell 166, 63–76 (2016).

14. Chao, T., Wang, H. & Ho, P.-C. Mitochondrial Control and Guidance of Cellular Activities of T Cells. Front. Immunol. 8 (2017).

15. Schumann, J. et al. The Mitochondrial Protein TCAIM Regulates Activation of T Cells and Thereby Promotes Tolerance Induction of Allogeneic Transplants. Am. J. Transplant. 14, 2723–2735 (2014).

16. Keeren, K. et al. Expression of Tolerance Associated Gene-1, a Mitochondrial Protein Inhibiting T Cell Activation, Can Be Used to Predict Response to Immune Modulating Therapies. J. Immunol. 183, 4077–4087 (2009).

17. Klein-Hessling, S. et al. NFATc1 controls the cytotoxicity of CD8(+) T cells. Nature Communications 8 (2017).

18. Hogan, P.G., Chen, L., Nardone, J. & Rao, A. Transcriptional regulation by calcium, calcineurin, and NFAT. Genes Dev. 17, 2205–2232 (2003).

19. Dimeloe, S., Burgener, A.V., Grahlert, J. & Hess, C. T-cell metabolism governing activation, proliferation and differentiation; a modular view. Immunology 150, 35–44 (2017).

20. Zorova, L.D. et al. Functional Significance of the Mitochondrial Membrane Potential. Biochem Mosc Suppl S 12, 20–26 (2018).

21. Phan, A.T. & Goldrath, A.W. Hypoxia-inducible factors regulate T cell metabolism and function. Mol. Immunol. 68, 527–535 (2015).

22. Pålsson-McDermott, E.M. & O’Neill, L.A.J. Targeting immunometabolism as an anti- inflammatory strategy. Cell Res. 30, 300–314 (2020).

23. Isaacs, J.S. et al. HIF overexpression correlates with biallelic loss of fumarate hydratase in renal cancer: Novel role of fumarate in regulation of HIF stability. Cancer Cell 8, 143–153 (2005).

24. Fischer, M. et al. Early effector maturation of naïve human CD8+ T cells requires mitochondrial biogenesis. Eur. J. Immunol. 48, 1632–1643 (2018).

25. Konjar, Š. et al. Mitochondria maintain controlled activation state of epithelial-resident T lymphocytes. Science Immunology 3, eaan2543 (2018).

26. López-Crisosto, C. et al. ER-to-mitochondria miscommunication and metabolic diseases. Biochimica et Biophysica Acta (BBA) - Molecular Basis of Disease 1852, 2096–2105 (2015).

27. Martinvalet, D. The role of the mitochondria and the endoplasmic reticulum contact sites in the development of the immune responses. Cell Death Dis. 9, 336 (2018).

28. Modi, S. et al. Miro clusters regulate ER-mitochondria contact sites and link cristae organization to the mitochondrial transport machinery. Nature Communications 10 (2019).

29. von der Malsburg, K. et al. Dual Role of Mitofilin in Mitochondrial Membrane Organization and Protein Biogenesis. Dev. Cell 21, 694–707 (2011).

30. Chen, H.W., Heiniger, H.J. & Kandutsch, A.A. Relationship between Sterol Synthesis and DNA-Synthesis in Phytohemagglutinin-Stimulated Mouse Lymphocytes. Proc. Natl. Acad. Sci. U. S. A. 72, 1950–1954 (1975).

31. Chakrabarti, R. & Engleman, E.G. Interrelationships between Mevalonate Metabolism and the Mitogenic Signaling Pathway in Lymphocyte-T Proliferation. J. Biol. Chem. 266, 12216–12222 (1991).

32. Kidani, Y. et al. Sterol regulatory element-binding proteins are essential for the metabolic programming of effector T cells and adaptive immunity. Nat. Immunol. 14, 489-+ (2013).

33. Yang, W. et al. Potentiating the antitumour response of CD8(+) T cells by modulating cholesterol metabolism. Nature 531, 651-+ (2016).

34. Sala-Vila, A. et al. Interplay between hepatic mitochondria-associated membranes, lipid metabolism and caveolin-1 in mice. Sci. Rep. 6 (2016).

35. Csordas, G. et al. Imaging Interorganelle Contacts and Local Calcium Dynamics at the ER-Mitochondrial Interface. Mol. Cell 39, 121–132 (2010).

36. De Vos, K.J. et al. VAPB interacts with the mitochondrial protein PTPIP51 to regulate calcium homeostasis. Hum. Mol. Genet. 21, 1299–1311 (2012).

37. Rizzuto, R. et al. Close contacts with the endoplasmic reticulum as determinants of mitochondrial Ca2+ responses. Science 280, 1763–1766 (1998).

38. Szabadkai, G. et al. Chaperone-mediated coupling of endoplasmic reticulum and mitochondrial Ca2+ channels. J. Cell Biol. 175, 901–911 (2006).

39. Anavi, S., Hahn-Obercyger, M., Madar, Z. & Tirosh, O. Mechanism for HIF-1 activation by cholesterol under normoxia: A redox signaling pathway for liver damage. Free Radic. Biol. Med. 71, 61–69 (2014).

40. Daley, S.R. et al. Rasgrp1 mutation increases naive T-cell CD44 expression and drives mTOR-dependent accumulation of Helios(+) T cells and autoantibodies. Elife 2 (2013).

41. Dodd, K.M., Yang, J., Shen, M.H., Sampson, J.R. & Tee, A.R. mTORC1 drives HIF-1 alpha and VEGF-A signalling via multiple mechanisms involving 4E-BP1, S6K1 and STAT3. Oncogene 34, 2239-2250 (2015).

42. Eid, W. et al. mTORC1 activates SREBP-2 by suppressing cholesterol trafficking to lysosomes in mammalian cells. Proc. Natl. Acad. Sci. U. S. A. 114, 7999–8004 (2017).

43. Porstmann, T. et al. SREBP activity is regulated by mTORC1 and contributes to Akt- dependent cell growth. Cell Metab. 8, 224–236 (2008).

44. Yang, K. et al. T Cell Exit from Quiescence and Differentiation into Th2 Cells Depend on Raptor-mTORC1-Mediated Metabolic Reprogramming. Immunity 39, 1043–1056 (2013).

45. Schieke, S.M. et al. The mammalian target of rapamycin (mTOR) pathway regulates mitochondrial oxygen consumption and oxidative capacity. J. Biol. Chem. 281, 27643–27652 (2006).

46. Head, S.A. et al. Antifungal drug itraconazole targets VDAC1 to modulate the AMPK/mTOR signaling axis in endothelial cells. Proc. Natl. Acad. Sci. U. S. A. 112, E7276–E7285 (2015).

47. Camara, A.K.S., Zhou, Y.F., Wen, P.C., Tajkhorshid, E. & Kwok, W.M. Mitochondrial VDAC1: A Key Gatekeeper as Potential Therapeutic Target. Front. Physiol. 8 (2017).

48. Ramanathan, A. & Schreiber, S.L. Direct control of mitochondrial function by mTOR. Proc. Natl. Acad. Sci. U. S. A. 106, 22229–22232 (2009).

49. Sood, A. et al. A Mitofusin-2-dependent inactivating cleavage of Opa1 links changes in mitochondria cristae and ER contacts in the postprandial liver. Proc. Natl. Acad. Sci. U. S. A. 111, 16017–16022 (2014).

50. Bantug, G.R. et al. Mitochondria-Endoplasmic Reticulum Contact Sites Function as Immunometabolic Hubs that Orchestrate the Rapid Recall Response of Memory CD8^+^ T Cells. Immunity 48, 542–555.e546 (2018).

## References Methods

51. Ocker, L. et al. Hypericin and its radio iodinated derivatives - A novel combined approach for the treatment of pediatric alveolar rhabdomyosarcoma cells in vitro. Photodiagnosis Photodyn. Ther. 29 (2020).

52. Kopka, J., et al.: the Golm Metabolome Database. Bioinformatics 21, 1635–1638 (2005).

53. Jaeger, C., Hoffmann, F., Schmitt, C.A. & Lisec, J. Automated Annotation and Evaluation of In-Source Mass Spectra in GC/Atmospheric Pressure Chemical Ionization-MS-Based Metabolomics. Anal. Chem. 88, 9386–9390 (2016).

54. Jaeger, C. & Lisec, J. Statistical and Multivariate Analysis of MS-Based Plant Metabolomics Data. Methods Mol. Biol. 1778, 285–296 (2018).

55. Schubert, M., Lindgreen, S. & Orlando, L. AdapterRemoval v2: rapid adapter trimming, identification, and read merging. BMC Res. Notes 9, 88 (2016).

56. Kim, D. et al. TopHat2: accurate alignment of transcriptomes in the presence of insertions, deletions and gene fusions. Genome Biol. 14, R36 (2013).

57. Langmead, B. & Salzberg, S.L. Fast gapped-read alignment with Bowtie 2. Nat Methods 9, 357–359 (2012).

58. Liao, Y., Smyth, G.K. & Shi, W. The R package Rsubread is easier, faster, cheaper and better for alignment and quantification of RNA sequencing reads. Nucleic Acids Res. 47 (2019).

59. Love, M.I., Huber, W. & Anders, S. Moderated estimation of fold change and dispersion for RNA-seq data with DESeq2. Genome Biol. 15 (2014).

60. Fresno, C. & Fernandez, E.A. RDAVIDWebService: a versatile R interface to DAVID. Bioinformatics 29, 2810–2811 (2013).

61. Yu, G.C., Wang, L.G., Han, Y.Y. & He, Q.Y. clusterProfiler: an R Package for Comparing Biological Themes Among Gene Clusters. Omics-a Journal of Integrative Biology 16, 284–287 (2012).

62. Walter, W., Sanchez-Cabo, F. & Ricote, M. GOplot: an R package for visually combining expression data with functional analysis. Bioinformatics 31, 2912–2914 (2015).

63. Thiele, I. et al. A community-driven global reconstruction of human metabolism. Nat. Biotechnol. 31, 419-+ (2013).

64. Lisec, J., Hoffmann, F., Schmitt, C. & Jaeger, C. Extending the Dynamic Range in Metabolomics Experiments by Automatic Correction of Peaks Exceeding the Detection Limit. Anal. Chem. 88, 7487–7492 (2016).

65. Karbowski, M. et al. Quantitation of mitochondrial dynamics by photolabeling of individual organelles shows that mitochondrial fusion is blocked during the Bax activation phase of apoptosis. J. Cell Biol. 164, 493–499 (2004).

66. Lehmann, A., Niewienda, A., Jechow, K., Janek, K. & Enenkel, C. Ecm29 Fulfils Quality Control Functions in Proteasome Assembly. Mol. Cell 38, 879–888 (2010).

67. Cox, J. et al. Accurate Proteome-wide Label-free Quantification by Delayed Normalization and Maximal Peptide Ratio Extraction, Termed MaxLFQ. Mol. Cell. Proteomics 13, 2513–2526 (2014).

68. Heinemeyer, T. et al. Databases on transcriptional regulation: TRANSFAG, TRRD and COMPEL. Nucleic Acids Res. 26, 362–367 (1998).

